# PRC1 and CTCF-Mediated Transition from Poised to Active Chromatin Loops Drives Bivalent Gene Activation

**DOI:** 10.1101/2024.11.13.623456

**Authors:** Aflah Hanafiah, Zhuangzhuang Geng, Tingting Liu, Yen Teng Tai, Wenjie Cai, Qiang Wang, Neil Christensen, Yan Liu, Feng Yue, Zhonghua Gao

## Abstract

Polycomb Repressive Complex 1 (PRC1) and CCCTC-binding factor (CTCF) are critical regulators of 3D chromatin architecture that influence cellular transcriptional programs. Spatial chromatin structures comprise conserved compartments, topologically associating domains (TADs), and dynamic, cell-type-specific chromatin loops. Although the role of CTCF in chromatin organization is well-known, the involvement of PRC1 is less understood. In this study, we identified an unexpected, essential role for the canonical Pcgf2-containing PRC1 complex (cPRC1.2), a known transcriptional repressor, in activating bivalent genes during differentiation. Our Hi-C analysis revealed that cPRC1.2 forms chromatin loops at bivalent promoters, rendering them silent yet poised for activation. Using mouse embryonic stem cells (ESCs) with CRISPR/Cas9-mediated gene editing, we found that the loss of Pcgf2, though not affecting the global level of H2AK119ub1, disrupts these cPRC1.2 loops in ESCs and impairs the transcriptional induction of crucial target genes necessary for neuronal differentiation. Furthermore, we identified CTCF enrichment at cPRC1.2 loop anchors and at Polycomb group (PcG) bodies, nuclear foci with concentrated PRC1 and its tethered chromatin domains, suggesting that PRC1 and CTCF cooperatively shape chromatin loop structures. Through virtual 4C and other genomic analyses, we discovered that establishing neuronal progenitor cell (NPC) identity involves a switch from cPRC1.2-mediated chromatin loops to CTCF-mediated active loops, enabling the expression of critical lineage-specific factors. This study uncovers a novel mechanism by which pre-formed PRC1 and CTCF loops at lineage-specific genes maintain a poised state for subsequent gene activation, advancing our understanding of the role of chromatin architecture in controlling cell fate transitions.

## Introduction

Epigenetic modulators, such as PcG proteins and CTCF, are central to regulating cellular transcriptional programs by shaping 3D chromatin architecture. Technological advances in high-throughput chromatin conformation capture techniques have led to the discovery of various levels of spatial organization within chromatin, including chromatin compartments, TADs, and long-range chromatin loops^1,2^. Unlike chromatin compartments and TADs^3,4^, which are highly conserved across different cell types, chromatin loops are dynamic and cell-specific^5^. CTCF plays a pivotal role in forming chromatin loops by coordinating with cohesin complexes^5,6^. Recent studies have shown that other proteins, including PcG proteins, establish chromatin loops via distinct mechanisms^7–13^. However, the interplay between PcG proteins and CTCF in regulating 3D chromatin architecture and, consequently, cell type-dependent transcriptomes, remains not fully understood.

PcG proteins are critical epigenetic regulators that modulate chromatin structure to maintain gene silencing, essential for stem cell maintenance and differentiation^14–17^. PcG proteins form two major protein complexes: Polycomb Repressive Complex 1 (PRC1) and 2 (PRC2)^14^. PRC1 and PRC2 catalyze two repressive chromatin modifications: mono-ubiquitination of histone H2A at lysine-119 (H2AK119ub1) and mono-, di- and tri-methylation of histone H3 at lysine-27 (H3K27me1/2/3), respectively^14^. Previously, we determined the composition of the mammalian PRC1 complexes, identifying six groups (PRC1.1-1.6) based on the exclusive association of one of the six Polycomb group RING fingers (PCGF1-6)^18^. Among these, PRC1.2 and PRC1.4 include the canonical PRC1 (cPRC1) complexes, initially isolated in *Drosophila*, and have homologous associated factors, including Really Interesting New Gene 1A/B (RING1A/B), Chromodomain proteins (CBX2/4/6/7/8) and Polyhomeotic homologs (PHC1/2/3)^18^.

Genome-wide analyses of gene targets in ESCs revealed the localization of cPRC1 complexes at regulatory sites of many developmental transcription factors (TF)^19–21^. Interestingly, subsequent studies discovered that H3K27me3, an inhibitory histone mark, and H3K4me3, an active mark, simultaneously bind promoters of these TFs^22^. The bivalent nature of these genes, due to the enrichment and enzymatic action of PRC2 for H3K27me3 and Mixed-Lineage Leukemia (MLL) for H3K4me3, maintains them at a silent yet poised state in ESCs^22^. Upon lineage-specific differentiation, bivalent genes critical for that particular lineage undergo rapid activation, and those for other lineages remain repressed^23^. It has been shown that cPRC1 complexes, including cPRC1.2 and cPRC1.4, are also present at bivalent promoters through the interaction with their CBX subunits to H3K27me3^24–26^. However, it remains unclear how cPRC1 complexes regulate the plasticity of bivalent gene expression during differentiation.

PRC1-mediated H2AK119ub1 is a hallmark of silenced chromatin^14^. While most PCGF proteins enhance the E3 ubiquitin ligase activity of RING1B for H2AK119ub1 deposition, PCGF2 and PCGF4 are exceptions to this pattern^27^. Removing Pcgf2 and Pcgf4 in mouse ESCs does not alter the overall levels of H2AK119ub1^28^, suggesting that the cPRC1 complexes may regulate gene transcription through mechanisms independent of H2AK119ub1. Indeed, cPRC1 components have been shown to mediate long-range chromatin interactions in mammalian cells, and disruption of multiple cPRC1 complex components leads to the loss of these loops^7–13^. Microscopy analysis further demonstrates that distal PRC1 target genes are localized within PcG bodies^29,30^—nuclear foci enriched with PcG proteins^31–37^—which contribute to the repression of their target genes, underscoring the regulatory role of PRC1-mediated long-range chromatin in gene expression.

This study aims to understand how the PRC1 complex regulates long-range chromatin interactions to control gene expression during cell fate determination. Using CRISPR/Cas9-mediated gene editing, we deleted Pcgf2 in ESCs. We found that Pcgf2 absence unexpectedly compromises the activation of PRC1-targeted bivalent genes upon differentiation, in contrast to its traditional role as a transcriptional repressor. Mechanistically, through various genomic analyses, we demonstrated that Pcgf2 is required to form a subset of chromatin loops that target bivalent promoters. These promoters, marked by H3K4me3 and H3K27me3, are also bound by CTCF. Although previous studies and our immunoprecipitation experiments showed no direct physical interactions between PRC1 and CTCF, immunofluorescence analysis revealed their colocalization at PcG bodies, indicating cooperation between PRC1 and CTCF in regulating high-order chromatin interactions. Importantly, our virtual 4C analysis showed that as ESCs differentiate into NPCs, PRC1 loops gradually diminish, and concomitantly, new chromatin loops mediated by CTCF emerge, many originating from former cPRC1.2 loop sites. Moreover, Pcgf2 deletion abolishes the CTCF-mediated active loops upon differentiation. Our findings uncover a previously unrecognized mechanism by which PRC1 and CTCF coordinate chromatin loop reorganization, a process essential for proper cell fate transition.

## Results

### 1. Pcgf2 regulates neuronal differentiation independently of H2AK119ub1

To investigate the roles of PRC1 complexes in regulating chromatin architecture during cell fate transition, we generated knockout ESC lines for *Pcgf2* (*Pcgf2^-/-^*) and *Pcgf4* (*Pcgf4^-/-^*) via CRISPR/Cas9-mediated gene editing (**Fig. S1a-b**). Genotyping PCR and immunoblotting confirmed the successful deletion of Pcgf2 and Pcgf4 (**Fig. S1c-f**). Using a previously established *in vitro* neuronal differentiation protocol (**Fig. S1g**)^38,39^, we found that Pcgf2, but not Pcgf4, was required for neuronal differentiation, as evidenced by immunofluorescence (IF) assay with a neurofilament (Nfm) antibody showing the defective dendritic growth in *Pcgf2^-/-^* neurons compared with wild type (*WT*) and *Pcgf4^-/-^* cells (**Fig. S1h**). The failed neuronal differentiation in *Pcgf2^-/-^* cells was also apparent at the NPC stage, with reduced expression of NPC marker genes, *Pax6* and *Neurod1*, as measured by RT-qPCR (**Fig. S1i**).

We then sought to determine whether the compromised neuronal differentiation in *Pcgf2^-/-^* cells was due to an impact on ESC self-renewal. Through alkaline phosphatase assay, we demonstrated that Pcgf2 deletion did not affect the ESC pluripotency in two independent *Pcgf2^-/-^*ESC lines, compared with *WT* (**Fig. S2a**). Similarly, there was no noticeable difference in cell proliferation in these *Pcgf2^-/-^* ESC lines, compared with *WT*, via MTT assay (**Fig. S2b**). Morphologically, when *Pcgf2^-/-^* ESCs were induced for NPC differentiation, we found no dramatic change in embryonic body (EB) shape but a slight reduction in EB size (**Fig. S2c-d**). Immunoblotting and RT-qPCR analysis showed that Pcgf2 deletion does not affect *Nanog*, a pluripotency marker, while reducing the expression of NPC marker genes, such as *Nes*, *Pax6*, and *Neurod1* (**Fig. S2e-f**). These data indicate that Pcgf2 deletion has no significant effect on ESC self-renewal and proliferation. Next, to investigate how the loss of Pcgf2 affects gene transcription during NPC differentiation, we performed RNA-seq analysis on *WT* and *Pcgf2^-/-^* cells across the ESC, EB, and NPC stages. We found that a group of genes was induced in *WT* NPCs but remained silent in *Pcgf2^-/-^* NPCs (**Fig. 1a**). Gene ontology (GO) analysis revealed that genes failed to be induced in *Pcgf2^-/-^* NPCs are primarily involved in cell differentiation and neuronal identity, including “pattern specification”, “axonogenesis”, “regionalization”, “neuron projection guidance”, and “axon guidance” (**Fig. 1b**).

**Figure 1.**
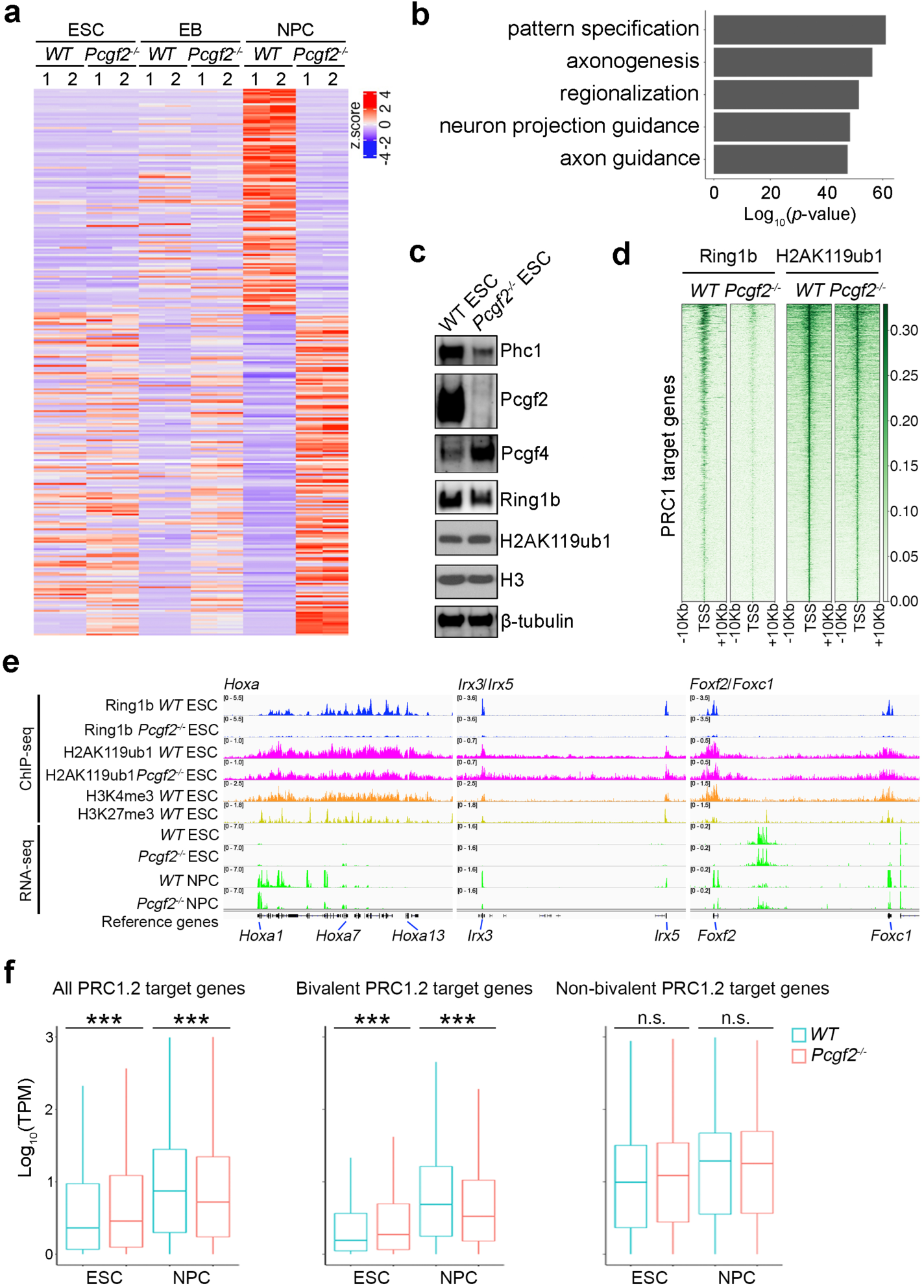
Pcgf2 is required for neuronal differentiation. **a**, Heat-map of differentially expressed genes (DEGs) in duplicated *WT* and *Pcgf2^-/-^* NPCs, clustered and sorted based on transcript z-score. A total of 2,263 upregulated and 1,576 downregulated genes were identified. **b**, Gene ontology (GO) analysis for downregulated genes in *Pcgf2^-/-^* NPCs from (**a**). **c**, Immunoblotting of cPRC1 components (Phc1, Pcgf2, Pcgf4, and Ring1b) and H2AK119ub1 in *WT* and *Pcgf2^-/-^* ESCs. **d**, ChIP-seq heatmaps showing H2AK119ub1 and Ring1b enrichment in *WT* and *Pcgf2^-/-^* ESCs at Ring1b (PRC1) target loci. **e**, ChIP-seq tracks for Ring1b, H2AK119ub1, H3K4me3, and H3K27me3 in *WT* and *Pcgf2^-/-^* ESCs and RNA-seq tracks for *WT* and *Pcgf2^-/-^* ESCs and NPCs at PRC1 target loci (*Hoxa*, *Irx3*/*Irx5*, and *Foxf2*/*Foxc1*). **f**, Box plots of expression levels for all PRC1.2 target genes (5,334), bivalent PRC1.2 target genes (3,284), and non-bivalent PRC1.2 target genes (2,050). \*\*\**p*<0.001; n.s., not significant.

To examine whether the observed transcriptomic changes by Pcgf2 deletion were caused by the compromised integrity of the PRC1 complex, we analyzed protein levels of PRC1 components by immunoblotting. Pcgf2 loss reduced the level of Phc1, a cPRC1 component (**Fig. 1c**), indicating that the Pcgf2 deletion caused the reduced stability of the cPRC1.2 complex. Ring1b, the E3 ligase for H2AK119ub1 and the common component of all PRC1 complexes, showed only a minor reduction in total protein levels in *Pcgf2^-/-^* cells (**Fig. 1c**), yet its genomic enrichment was decreased (**Fig. 1d**). However, consistent with previous findings^28^, global H2AK119ub1 level was not substantially affected in *Pcgf2^-/-^* cells, evidenced by both immunoblotting and ChIP-seq analysis (**Fig. 1c-d**), suggesting a potential H2AK119ub1-independent mechanism mediating the effect by Pcgf2 loss. Interestingly, Pcgf2 deletion resulted in Pcgf4 overexpression, possibly compensating for Pcgf2 loss (**Fig. 1c**) but unable to rescue neuronal differentiation defects caused by Pcgf2 knockout (**Fig. 1a-b**). Altogether, our results reveal an H2AK119ub1-independent role for PRC1.2 in regulating NPC differentiation.

### 2. Pcgf2 deletion compromises activation of PRC1.2-targeted bivalent genes upon neuronal differentiation

Previous studies have shown that PRC1 complexes localize at bivalent promoters^24,40^. We sought to investigate if PRC1.2 plays a role in regulating bivalent gene expression during neuronal differentiation. We first identified bivalent loci simultaneously marked by the H3K27me3 and H3K4me3. We then cross-referenced these loci with Ring1b and Pcgf2 binding sites on the genome, identifying a total of 3,284 PRC1.2-targeted bivalent promoters. Three examples are shown for the *Hoxa*, *Irx3/Irx5*, and *Foxf2/Foxc1* loci (**Fig. 1e**). These PRC1.2 target genes are kept silent in *WT* ESCs, and Pcgf2 deletion only mildly increases their expression (**Fig. 1e**), which is consistent with a previous report^28^. When neuronal differentiation was initiated, we observed dramatic induction of these genes in *WT* NPCs (**Fig. 1e**), in keeping with their roles in neurodevelopment as previously described^41–43^. The Pcgf2 deletion inhibited the activation of PRC1.2 target genes (**Fig. 1e**), which is somewhat surprising given the traditional roles of PRC1 as transcriptional repressors. Globally, PRC1.2 target genes tend to be less activated in *Pcgf2^-/-^*NPCs compared to *WT* NPCs (**Fig. 1f**). Interestingly, when we divided PRC1.2 target genes based on their bivalency status, we found a great reduction in transcriptional activation of the bivalent PRC1.2 target genes in *Pcgf2^-/-^* NPCs, but there was no noticeable change in the non-bivalent PRC1.2 target genes (**Fig. 1f**). Our results suggest that the PRC1.2 complex specifically target bivalent genes for their activation upon differentiation.

### 3. cPRC1.2 mediates long-range chromatin interactions

Previous studies have demonstrated that PRC1 complexes regulate long-range chromatin interactions independently of Ring1b catalytic activity toward H2AK119ub1^44^. Furthermore, Phc1, a component essential for cPRC1 oligomerization^45,46^, has been shown to influence long-range chromatin interactions at selective loci in ESCs^10^. Given that Pcgf2 deletion had little effect on H2AK119ub1 but notably reduced Phc1 level (**Fig. 1c**), we speculated that the observed dysregulation of bivalent genes in *Pcgf2^-/-^* NPCs was mediated by the cPRC1.2 complex through long-range chromatin interactions. To test this hypothesis, we performed Hi-C analysis in *WT* and *Pcgf2^-/-^* ESC samples. Deep-sequenced raw data were mapped to the mouse genome GRCm38 to detect ligation junctions (**Table S1**). Our bioinformatic analysis identified 3,912 chromatin loops in *WT* ESCs; among them, 458 are not present in *Pcgf2^-/-^* ESCs, which we designated as PRC1.2 dependent loops (**Fig. S3a**). **Fig. 2a** shows three selected examples, including *Hoxb*, *Wt1*/*Pax6*, and *Irx3*/*Irx5* loci, with substantially reduced contact intensities in *Pcgf2^-/-^* Hi-C matrix compared to *WT*. Pcgf2 and Ring1b, two core components of the cPRC1.2 complex, are enriched at the genomic anchors of these loops (**Fig. 2a**). The *Hoxb* and *Pax6* loci have been shown in previous studies as targeted by PRC1 loops^10^. We observed no significant change in A/B compartments and TADs in *Pcgf2^-/-^* ESCs compared to *WT* at these loci (**Fig. S3b-d**). **Table S2** shows a complete list of the genes targeted by PRC1.2 loops.

**Figure 2.**
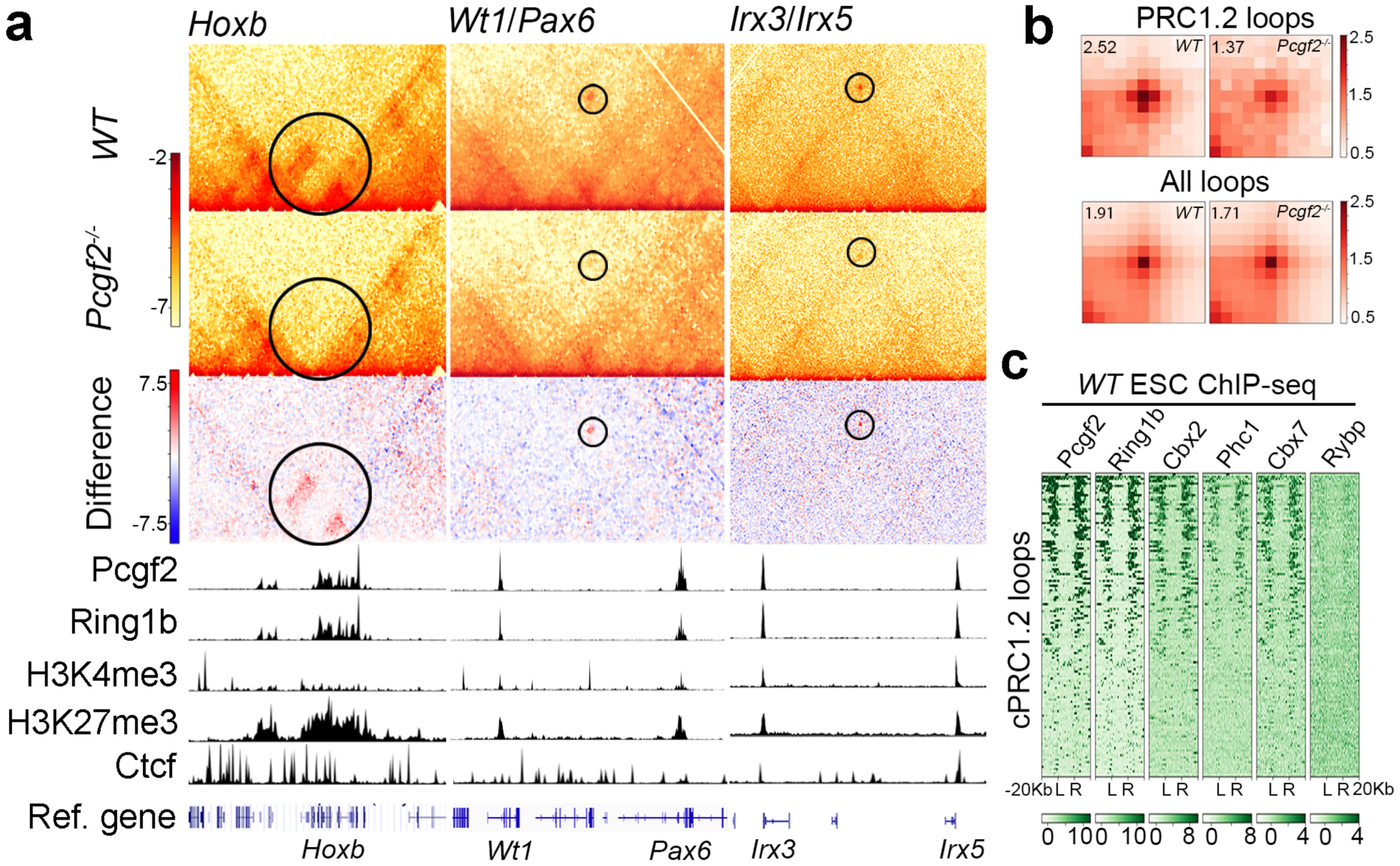
cPRC1.2 is required for chromatin loop formation. **a,** Example Hi-C matrices for *WT* and *Pcgf2^-/-^* ESCs at 5-kb resolution showing reduced contact intensities at *Hoxb*, *Wt1*/*Pax6*, and *Irx3*/*Irx5* loci in *Pcgf2^-/-^*ESCs. ChIP-seq tracks for Pcgf2, Ring1b, H3K4me3, H3K27me3, and Ctcf are displayed below the Hi-C matrices. **b,** Aggregate peak analysis (APA) of Hi-C for total chromatin loops and cPRC1.2 loops in *WT* and *Pcgf2^-/-^*ESCs at 10-kb resolution. The pileups are normalized to the average of the top-left and bottom-right corner pixels, and the value of the central pixels is displayed on the top-left side of the plot. **c**, ChIP-seq heatmaps show enrichment of cPRC1.2 components (Pcgf2, Cbx2, Phc1, and Cbx7) and lack of ncPRC1 component (Rybp) at cPRC1.2 loop anchors in *WT* ESCs. L and R denote left and right loop anchors, with 20 kb regions flanking each anchor.

Aggregate peak analysis (APA), which measures average loop strength, revealed a global reduction in PRC1.2-associated chromatin loops in *Pcgf2^-/-^*ESCs (**Fig. 2b**, upper panel). Although CTCF is localized to these loop anchoring regions (**Fig. 2a**), there is no noticeable difference in total chromatin loops between *WT* and *Pcgf2^-/-^* ESCs (**Fig. 2b**, lower panel), suggesting that the effect of Pcgf2 deletion is restricted to the selected loops targeted by PRC1.2. Interestingly, these PRC1.2 loops are often anchored at sites with enrichment of H3K4me3 and H3K27me3, suggesting a bivalent status of these regions (**Fig. 2a**). Further ChIP-seq analysis revealed that the cPRC1.2 components such as Pcgf2, Ring1b, Cbx2, Phc1, and Cbx7, are all enriched at loop anchors, but Rybp, an ncPRC1 component, is absent (**Fig. 2c**). In summary, our results demonstrate that the cPRC1.2 complex is responsible for the establishment of a specific set of chromatin loops.

### 4. cPRC1.2 loops target bivalent genes for activation upon neuronal differentiation

Our observation of bivalent histone marks at selected cPRC1.2-mediated chromatin loop anchors (**Fig. 2a**) suggests the potential involvement of these loops in regulating bivalent genes. Therefore, we conducted a further bioinformatic analysis to assess the genome-wide relationship between the cPRC1.2-mediated chromatin loops and bivalency. As shown in **Fig. 3a**, cPRC1.2-mediated chromatin loops are primarily anchored at transcription start sites (TSS). More importantly, cPRC1.2 loop anchors are found to be enriched with H3K4me3 and H3K27me3 modifications globally compared to the total loop regions identified (**Fig. 3b**). Our analysis showed that PRC1 loops exclusively target bivalent promoters, with a strikingly high percentage (94%) of cPRC1.2 loops-targeted promoters being bivalent, compared with 56% of all PRC1-targeted and 54% of non-loop forming PRC1-targeted promoters showing bivalency (**Fig. 3c**).

**Figure 3.**
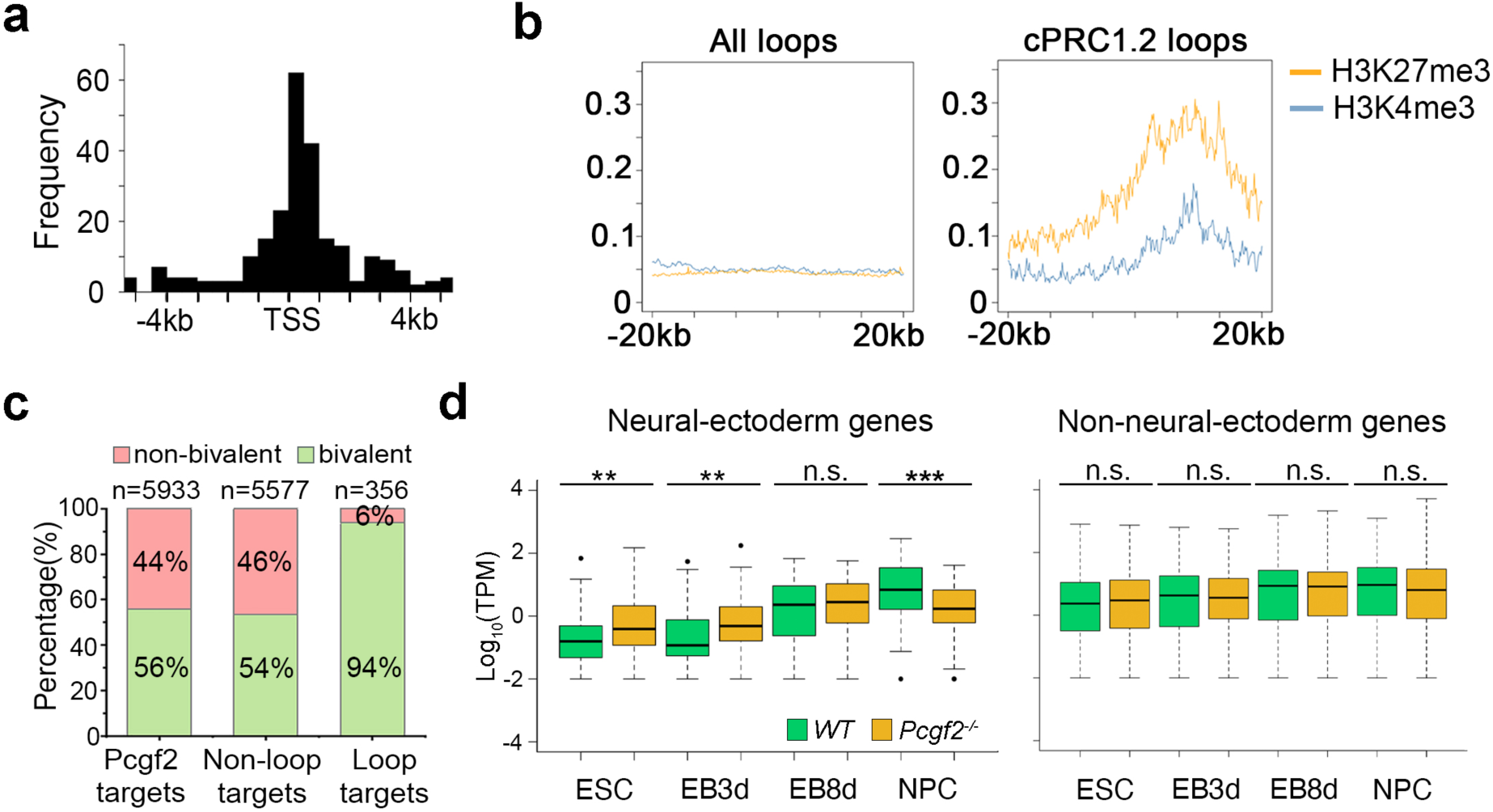
cPRC1.2 loops target bivalent promoters for gene activation upon differentiation. **a**, Distribution of Pcgf2-dependent loop regions relative to the nearest annotated transcription start site (TSS). The x-axis represents the distance to TSS. **b**, ChIP-seq signal showing H3K27me3 and H3K4me3 enrichment across all loop regions (left panel) and cPRC1.2 loop regions (right panel) in *WT* ESCs. **c**, Percentage of bivalent domains within total Pcgf2 targets, non-loop Pcgf2 targets, and cPRC1.2 loop targets. **d**, Box plots showing expression of neural-ectoderm genes (82) and non-neural-ectoderm genes (372) that are targeted by cPRC1.2 loops in *WT* and *Pcgf2^-/-^* cells across ESC, EB, and NPC stages. \*\**p*<0.01; *** *p*<0.001; n.s., not significant.

In our transcriptomic analysis, we discovered a surprisingly reduced expression of bivalent genes targeted by cPRC1.2 in *Pcgf2^-/-^*NPCs (**Fig. 1e-f**). Given the specific targeting of bivalent promoters by cPRC1.2 loops, we then examined how these chromatin loops may affect the activation of bivalent genes targeted by PRC1. Although no difference was found in averaging transcript per million (TPM) values for all PRC1 target genes and cPRC1.2 loop-targeted genes between *WT* and *Pcgf2^-/-^*NPCs (**Fig. S4a**), Pcgf2 deletion led to a significant reduction in TPM for neuro-ectodermal genes targeted by the cPRC1.2 loops in *Pcgf2^-/-^* NPCs (**Fig. 3d and Fig. S4b**). Interestingly, either non-neuro-ectodermal or meso-endodermal genes targeted by the cPRC1.2 loops showed no noticeable difference in their expression between *WT* and *Pcgf2^-/-^* NPCs (**Fig. 3d and Fig. S4a**), indicating that cPRC1.2 loops are selectively required for the activation of neuro-ectodermal genes in our neuronal differentiation model. When we omitted retinoic acid (RA) at day 4 and let EB differentiate without restriction into NPC in the same differentiation period (EB8d), the observed difference in expression of neuro-ectodermal genes targeted by cPRC1.2 loops was diminished (**Fig. 3d**). It is worth noticing that, at the ESC stage, the Pcgf2 deletion causes a de-repression of neuro-ectodermal genes targeted by cPRC1.2 loops but has no noticeable effect on non-neuro-ectodermal or meso-endodermal genes (**Fig. 3d and Fig. S4a**). Our results identify a novel role for the cPRC1.2 complex in activating neuro-ectodermal lineage genes through chromatin looping to promote NPC identity.

### 5. Phc1 SAM domain deletion in ESCs mimics Pcgf2 knockout effects

Pcgf2 is also associated with ncPRC1 components such as Rybp^18,47,48^. To further evaluate the specific contribution of cPRC1.2-mediated loops in controlling the expression of bivalent genes, we generated an ESC line with the deletion of Phc1 sterile alpha motif (*Phc1ΔSAM*) (**Fig. 4a, Fig. S5a**). Phc proteins are components specific to cPRC1^18^. Among all three Phc paralogs, Phc1 is the predominant form in ESCs^49^, and the *Phc1* deletion in ESCs disrupted selective chromatin loops^10^. It has been previously shown that the SAM domain of Phc mediates PRC1 oligomerization and PRC1 clustering in cells^45,46^. The *Phc1ΔSAM* ESC line has been confirmed by Sanger sequencing, genotyping PCR, and immunoblotting (**Fig. S5a-b, Fig. 4b**, note the smaller size of *Phc1ΔSAM*). The SAM domain deletion has no noticeable effect on global protein levels in PRC1 components, including Ring1b, Pcgf2, Rybp, and PRC1-mediated modification H2AK119ub1 (**Fig. 4b**).

**Figure 4.**
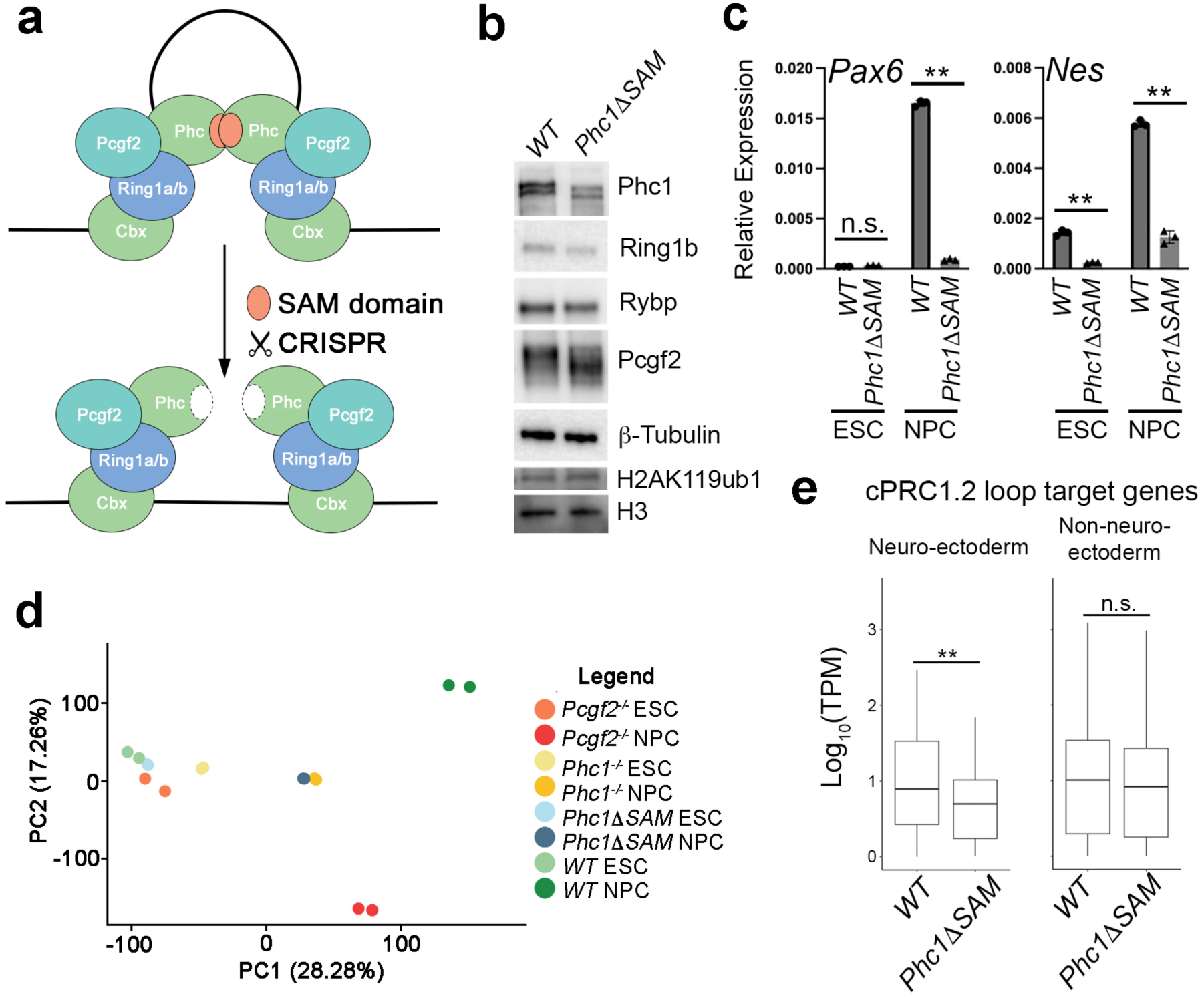
Deleting Phc1 SAM domains compromises target gene activation in NPCs. **a**, Schematic of Phc1 SAM domain deletion (*Phc1ΔSAM*) for disrupting cPRC1 oligomerization. **b**, Immunoblotting of Phc1, Ring1b, Rybp, Pcgf2, and H2AK119ub1 in *WT* and *Phc1ΔSAM* ESCs. Note the smaller size of the Phc1 band in the *Phc1ΔSAM* sample. **c**, RT-qPCR analysis showing failed induction of NPC markers (*Pax6* and *Nes*) in *WT* and *Phc1ΔSAM* NPCs. n = 3 for each sample. **d**, Principal component analysis (PCA) of RNA-seq data from *WT*, *Phc1^-/-^*, and *Phc1ΔSAM* ESCs and NPCs. **e**, Box plots of expression of neuro-ectodermal and non-neuro-ectodermal genes targeted by cPRC1.2 loops in *WT* and *Phc1ΔSAM* NPCs. \*\**p*<0.01; n.s., not significant.

To examine the impact of *Phc1ΔSAM* on neuronal gene activation, we differentiated the *Phc1ΔSAM* and *WT* ESCs into NPCs. Through RT-qPCR analysis, we found that disrupting the Phc1 SAM domain caused the failure of induction of NPC markers (*Pax6* and *Nes*) (**Fig. 4c**), suggesting a similar NPC differentiation defect as seen in *Pcgf2^-/-^* cells (**Fig. 1** and **Fig. S1**). Further transcriptomic analysis showed that many genes related to neuronal cell fate transition were downregulated in *Phc1ΔSAM* NPCs compared with *WT* (**Fig. S5c-d**). To further assess the transcriptomic changes caused by Pcgf2 and Phc1 disruption, we performed a Principal Component analysis (PCA). We also created a Phc1 total knockout (*Phc1^-/-^*) ESC cell line and then conducted RNA-seq for the comparison (**Fig. S5e-f**). As shown in **Fig. 4d**, all ESC samples, including *WT*, *Pcgf2^-/-^*, *Phc1^-/-^*, and *Phc1ΔSAM*, are clustered closely, indicating a minor effect on gene expression caused by cPRC1.2 disruption. However, NPC samples are scattered more distantly, with *WT* and *Pcgf2^-/-^* being the furthest apart from each other, and *Phc1^-/-^* and *Phc1ΔSAM* are placed relatively closer to *Pcgf2^-/-^* compared to *WT* (**Fig. 4d**). This may reflect the more dramatic impact of Pcgf2 deletion on both cPRC1 and ncPRC1 than Phc1 disruption with restricted effect on cPRC1. Our early analysis in *Pcgf2^-/-^* NPCs revealed a specific failure to induce neuro-ectodermal genes targeted by the cPRC1.2 loops (**Fig. 3d**). When we examined the impact of *Phc1ΔSAM* on PRC1.2 loop-targeted genes, we found that SAM deletion led to a similar observation, with a reduction in cPRC1.2 loop-targeted gene expression related to neuro-ectoderm but not the ones related to other lineages (**Fig. 4e**). In summary, by disrupting Phc1 through SAM domain deletion, our results further support the role of cPRC1.2-mediated loops for activation of genes critical for neuronal lineage differentiation.

### 6. CTCF is colocalized with cPRC1.2

Our Hi-C analysis identified CTCF enrichment at anchors of selected cPRC1.2 chromatin loops (**Fig. 2a**), raising whether CTCF cooperates with PRC1 in chromatin looping. Interestingly, a previous report has shown that CTCF binding is highly correlated with chromatin loops established by PRC2^50^, which catalyzes H3K27me3 as docking sites for cPRC1 binding^25,26^. To examine the global chromatin distribution of CTCF in relation to cPRC1.2 loops, we calculated the binding intensities of PRC1.2 and CTCF at cPRC1.2 loop anchors. As a result, both Ring1b and Pcgf2 show strong genomic enrichment at these loci (**Fig. 5a**). CTCF is enriched at the loop anchors, albeit with less intensity and with sustained distribution in the neighboring regions flanking its peaks (**Fig. 5a**).

**Figure 5.**
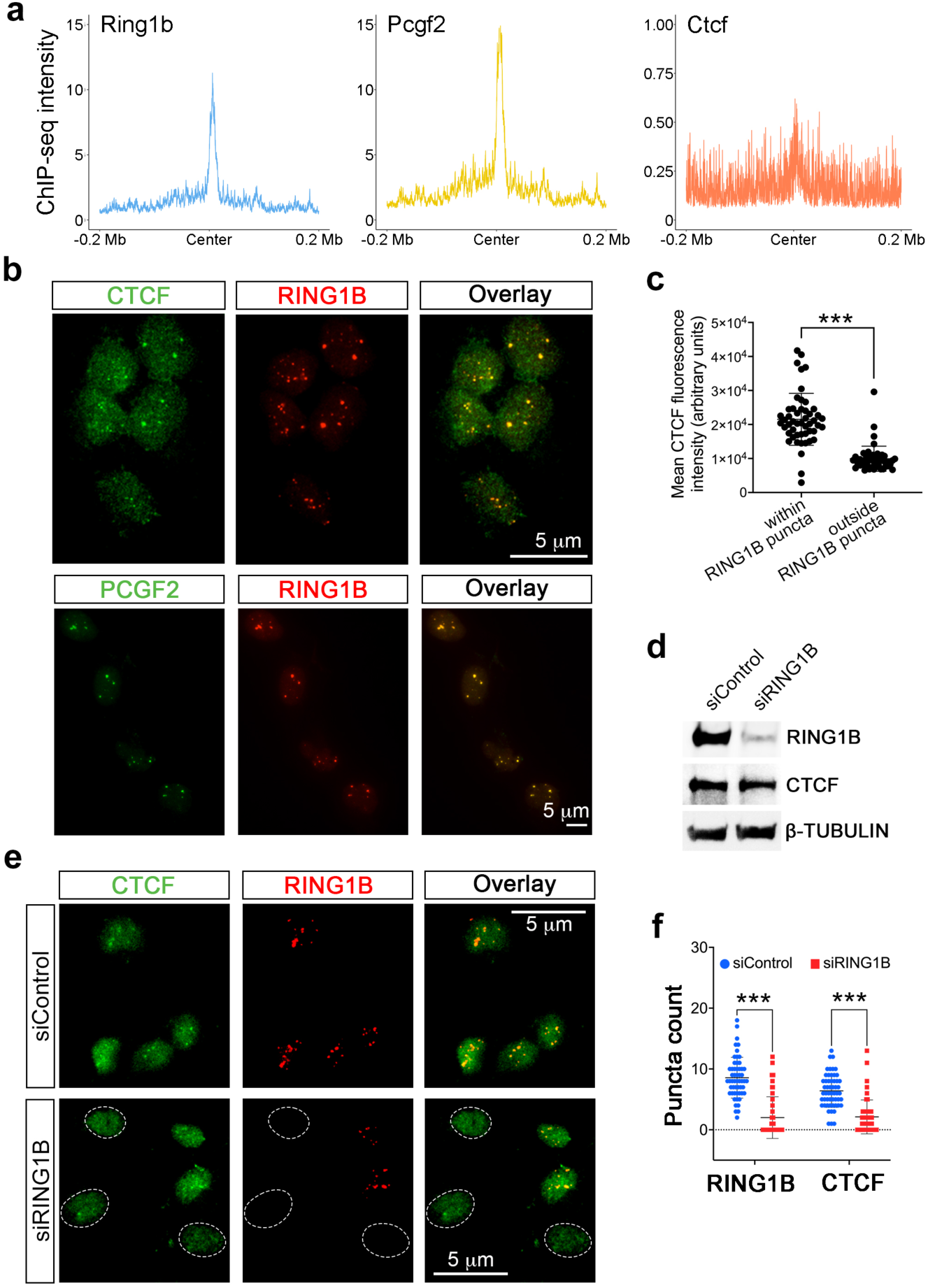
CTCF is colocalized at cPRC1.2 sites. **a**, ChIP-seq enrichment of Ring1b, Pcgf2, and CTCF at cPRC1.2 loop anchors with 0.2 Mb regions flanking each anchor in *WT* ESCs. **b,** Immunofluorescent images of U2OS cells co-stained for CTCF and RING1B (top panel) and PCGF2 and RING1B (bottom panel). **c**, Quantification of CTCF fluorescence within or outside RING1B puncta (n = 50). **d**, Immunoblotting of RING1B and CTCF in RING1B knock-down (siRING1B) and control (siControl) U2OS cells. **e**, Immunofluorescent images of control (siControl) and RING1B knock-down (siRING1B) U2OS cells showing a reduction in CTCF puncta with RING1B knockdown. White circular outlines indicate the nuclei boundaries for cells with successful knockdown. **f**, Dot plots showing the number of RING1B and CTCF puncta in siControl (n = 54) and siRING1B (n = 50) U2OS cells. \*\*\**p*<0.001.

Direct physical interactions between PRC1 and CTCF have not been detected through affinity purification studies in the past^18,19,51–56^. We did not observe their interaction by immunoprecipitation either (**Fig. S6a**). cPRC1 components are known to be concentrated in PcG bodies, nuclear structures that serve as hubs to tether distantly located PRC1 targeted loci to maintain their silencing^29,30^. Interestingly, CTCF has been found to be located at PcG bodies, but the functional importance remained unclear^57^. Therefore, to understand the relationship of CTCF with PRC1-associated PcG bodies, we performed IF studies in U2OS cells, which have been used previously for their ease of visualizing PcG bodies^32,33,36,37^. Strong CTCF punctate signals were detected within RING1B-positive PcG bodies (**Fig. 5b**, top panel). PCGF2 also showed a high level of colocalization with RING1B (**Fig. 5b**, bottom panel), indicating the presence of the cPRC1.2 complex at PcG bodies. We quantified the CTCF fluorescent intensity within or outside PcG bodies and found that CTCF was highly concentrated at RING1B-enriched PcG bodies (**Fig. 5c**).

CTCF may play a causative role in PcG body formation, or, inversely, its localization at PcG bodies depends on PRC1. To test these possibilities, we performed siRNA-mediated knockout-down analyses. In U2OS cells treated with siRNA for RING1B, we achieved a high level of RING1B silencing compared with cells treated with control siRNAs, demonstrated by immunoblotting analysis (**Fig. 5d**). The loss of RING1B expression in U2OS cells led to the disruption of PcG bodies shown by the lack of fluorescent RING1B and an accompanied disappearance of CTCF puncta, compared to control (**Fig. 5e**). Overall, we observed a reduction in nuclear puncta positive for both RING1B and CTCF upon RING1B knockdown (**Fig. 5f**). On the other hand, when we knocked down CTCF (**Fig. S6b**), we did not see a dramatic change in RING1B-positive foci (**Fig. S6c-d**), suggesting CTCF is not required for PcG body formation. Altogether, these results suggest a potential collaboration between cPRC1.2 and CTCF in proximity for regulating chromatin loops and gene activity.

### 7. Disruption of cPRC1.2 inhibits the formation of active loops mediated by CTCF upon neuronal differentiation

Given our observation of the localization of CTCF in cPRC1.2 loop anchors as well as in PcG bodies (**Fig. 5**), we hypothesized that CTCF may coordinate with cPRC1.2 to regulate the lineage-specific gene activation through chromatin looping. To investigate this, we performed a virtual 4C analysis on our Hi-C data from *WT* and *Pcgf2^-/-^* ESCs and previously published Hi-C data in *WT* ESC and NPCs^58^. We also generated Hi-C data in *Pcgf2^-/-^* NPCs to investigate the impact of the inactivation of the PRC1.2 complex. We first chose one loop anchor as the bait and then calculated its contact frequencies with its neighboring regions. At the *Foxf2/Foxc1* locus, we observed that the interaction strength of the previously identified cPRC1.2 loop was notably weakened at the NPC stage (**Fig. 6a**, top panel, purple-shaded peaks). Globally, cPRC1.2 loops that target the neuro-ectodermal genes often show a decreased loop strength upon differentiation into NPCs (**Fig. 6d**). In contrast, cPRC1.2 loops targeting non-neuro-ectodermal genes show no significant change (**Fig. 6d**). Interestingly, accompanying the weakening of cPRC1.2 loop at the *Foxf2/Foxc1* locus upon differentiation, the bait showed increased contact with adjacent genomic regions bound by CTCF but outside the cPRC1.2 loop (**Fig. 6a and 6c**, green-shaded areas), indicating the enhancement of CTCF-mediated chromatin loops. Furthermore, Pcgf2 deletion weakened these newly formed loops in *Pcgf2^-/-^*NPCs (**Fig. 6a**, bottom), which suggests that cPRC1.2 is required for the formation of subsequent CTCF loops.

**Figure 6.**
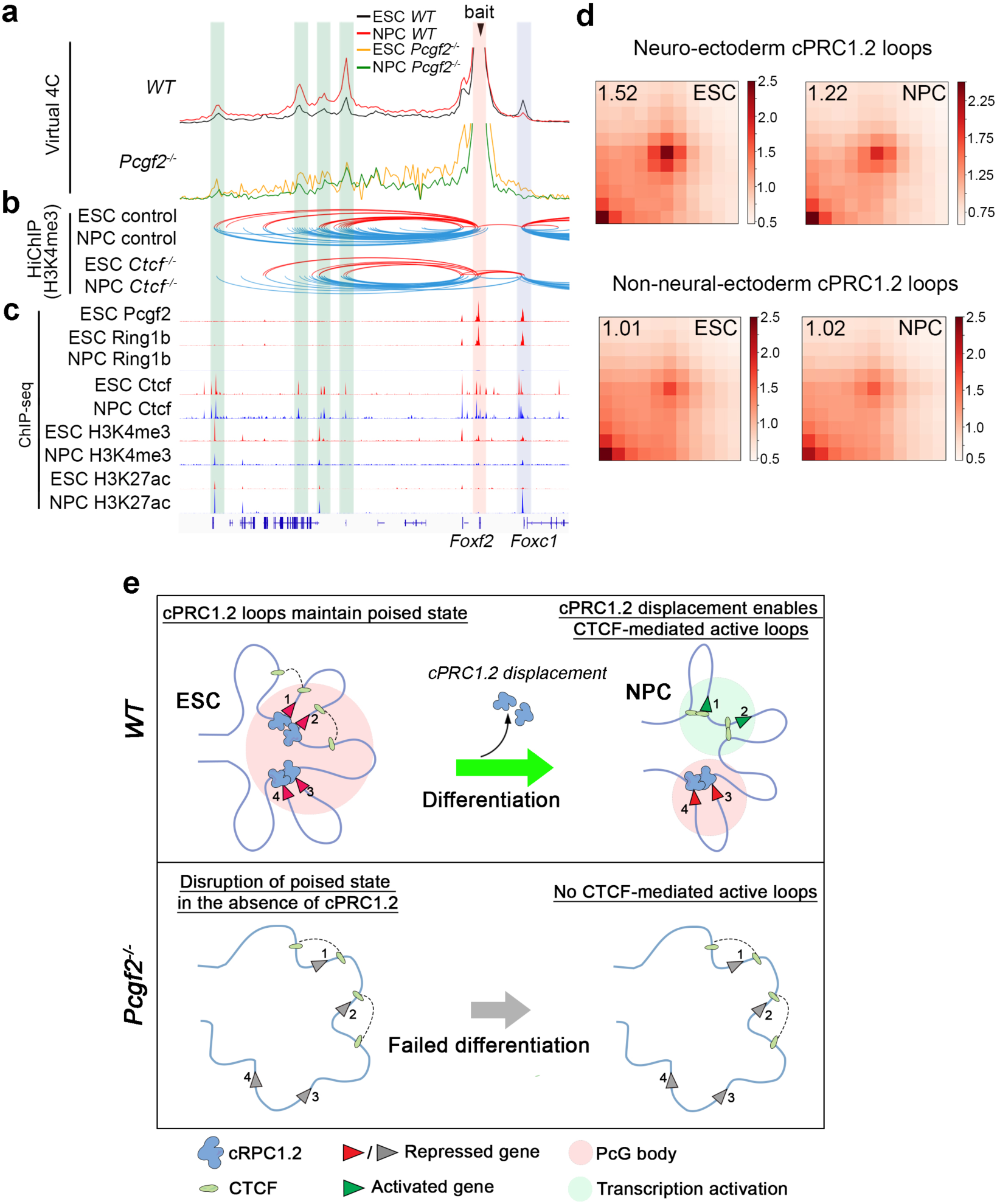
cPRC1 loops prime subsequent active CTCF loops during differentiation. **a,** Virtual 4C analysis of *Foxf2/Foxc1* locus using Hi-C data in *WT* and *Pcgf2^-/-^* ESCs and NPCs. The bait was set at the loop anchor at *Foxf2* (highlighted in red). Note the increased interactions in NPCs outside of the loop domain (highlighted in green) and the decreased interaction between *Foxf2* and *Foxc1* (highlighted in purple) in the *WT* cells. Interactions are normalized based on the peak height of the bait. **b**, HiChIP analysis for H3K4me3, from a previous study^46^, shows the increase of promoter-enhancer or promoter-promoter loops in *WT* NPCs compared with ESCs (top panel). These loops are weakened when Ctcf is deleted (bottom panel). These loops are well correlated with the Virtual 4C peaks. **c**, ChIP-seq tracks showing the enrichment for Pcgf2, Ring1B, Ctcf, H3K4me3, and H3K27ac in *WT* ESCs and NPCs. **d**, APA plots for cPRC1.2 loops targeting neuro-ectoderm genes (top) or non-neuro-endoderm genes (bottom) in *WT* ESCs and NPCs at 10-kb resolution. Normalized pileup values are displayed at the top-left. **e**, Schematic model illustrating the role of cPRC1.2 loops in regulating lineage-specific gene expression through establishing a primed state for subsequent CTCF-mediated loops during neuronal differentiation.

A previous study, through HiChIP analysis, using an H3K4me3 antibody, identified promoter-enhancer and promoter-promoter loops gained during NPC differentiation^59^. Furthermore, the formation of these loops requires the presence of CTCF and promotes gene activation^59^. To test whether the newly gained loops we observed in NPCs were previously identified CTCF-mediated active loops, we took advantage of the published HiChIP dataset and extracted the H3K4me3 loops surrounding our identified cPRC1.2 loop sites. As a result, we found an increase of H3K4me3 HiChIP contacts between the bait and the genomic regions, showing enhanced virtual 4C contacts (**Fig. 6b**, green-shaded areas). In addition, these H3K4me3 loops are dependent on CTCF since auxin-induced CTCF deletion (*Ctcf^-/-^*) led to reduced H3K4me3 HiChIP contacts (**Fig. 6b**, compare NPC control and NPC *Ctcf^-/-^*).

It has been suggested that the distal promoter may act as an enhancer to stimulate the gene activity downstream of the proximal promoter^60,61^. Therefore, we used the H3K4me3/H3K27ac ratio to gauge the status of the distal genomic elements looped to the proximal promoter targeted by cPRC1.2 loops. As seen in **Fig. 6c**, an increase in H3K27ac occupancy and a decrease in H3K4me3 level were observed at distal looped regions upon differentiation, suggesting the enhancer activity of the distal elements. Taken together, our results uncover a novel mechanism linking PRC1 and CTCF to orchestrate chromatin looping interactions for bivalent gene activation that drives neuronal differentiation (**Fig. 6e**).

## Discussion

Our study uncovers a novel mechanism through which PRC1, traditionally linked to gene repression, plays an unexpected role in activating essential developmental genes by collaborating with CTCF to reorganize chromatin topology. The cPRC1.2 complex forms poised chromatin loops at bivalent promoters of critical lineage-specific TFs in ESCs, and upon neuronal differentiation, cPRC1.2 loops dissolve with simultaneous enhancement of pre-formed CTCF-mediated active loops (**Fig. 6e**, top panel). This rearrangement of chromatin structure allows for the timely activation of key TFs for the transition to NPCs. The loss of cPRC1.2 leads to the disruption of PRC1 loops in ESCs, which compromises the formation of subsequent CTCF loops and ultimately prevents neuronal TF induction (**Fig. 6e**, bottom panel).

PRC1 complexes perform gene regulatory roles by maintaining a repressive chromatin environment^14–17^. Recent evidence suggests they can also function as transcriptional activators^62–67^. Despite the previous studies establishing the critical role of PRC1-mediated chromatin loops in transcription, whether they positively or negatively regulate gene activity remains controversial. Although earlier studies mostly align with a repressive role of PRC1 loops^8,68^, it has been later observed that PRC1-mediate promoter-enhancer loops are formed in both *Drosophila* eye-antennal imaginal discs and mouse brains, resulting in target gene activation^12,13^. Our proposed model may provide a means to reconcile these opposing observations. At the ESC stage, PRC1 loops target bivalent promoters to maintain their silencing while prime them for activation by subsequent topological rearrangement. Upon differentiation, PRC1 loops are weakened or disassembled to allow their targeted promoters to form CTCF-mediated loops with distal regulatory elements, leading to gene activation. This model is indeed consistent with the previous observation at the *Meis2* locus in the developing mouse brains, where a temporal transition of PRC1-mediated loop occurs between its promoter and a site within the 3’-region to a promoter-enhancer loop^12^. Interestingly, this study identified an intermediate tripartite loop formed by these elements^12^, although the molecular mechanism was unclear. We suspect that the colocalization of CTCF in PcG bodies may provide a looping mechanism to link the *Meis2* promoter to the corresponding enhancer. It will be interesting to test whether this is the case and extend further studies to elucidate how distinct types of loops dynamically regulate chromatin reorganization to activate key lineage-specific factors during development.

PcG bodies have long been observed in cells from both *Drosophila* and mammalian origins^29–37^. Recently, chromosomal phase separation has been proposed to interpret the formation of nuclear condensates, including various nuclear bodies such as PcG bodies, heterochromatin, and transcriptional condensates^69–74^. Such nuclear condensates may act as structural hubs for either active gene transcription or repression. In particular, PcG bodies have been shown to organize looped PRC-associated genes for their silencing^29,30^. Our findings of the co-localization of cPRC1.2 and CTCF in PcG bodies (**Fig. 5**) suggest that PcG bodies may also serve as a platform for priming genes for activation upon differentiation. Future studies will be needed to address how these two mechanisms may cooperate to achieve sophisticated gene regulation. Regarding the collaboration between CTCF and PRC1, another critical question remains: how is CTCF recruited to PcG bodies? We have shown a clear dependence of CTCF on the RING1B to be enriched in such nuclear bodies, but it is unlikely through direct interaction between CTCF and cPRC1.2. As previously reported, PRC2 is also localized to PcG bodies, so the reported interaction of PRC2 subunits with CTCF^75,76^ may assist the CTCF recruitment. Alternatively, an indirect mechanism may enhance CTCF retention on chromatin in the compacted PcG bodies.

Given the cell-type-dependent complexity of the PRC1 composition^18,49^, future effort will be needed to clarify whether the different cell or animal models used in previous studies may account for the differential regulation by cPRC1 loops on distinct genes. For example, given their nearly identical subunit makeups^18^, does cPRC1.4 have a similar effect on loops as cPRC1.2? Although our data suggest a predominant role for cPRC1.2 in regulating neuronal genes *in vitro*, we cannot rule out the possibility that cPRC1.4 or cPRC1.2 may be necessary for other lineages or even different neuronal cell types. Recently, it has been reported that patients harbor a missense mutation in the *Pcgf2* locus, resulting in a developmental syndrome called the Turnpenny-Fry syndrome. The developmental defects included brain tissue malformations accompanied by a varying degree of intellectual disability^77–79^. These reports provide an important rationale for studying cPRC1 functions in regulating chromatin architecture in the context of development in other lineages besides neuronal lineages.

## Acknowledgements

We thank the Genome Sciences core at Pennsylvania State University College of Medicine (RRID: SCR_021123), especially Dr. Sirisha Pochareddy, Dr. Yuka Imamura, and their team members, for assisting in library construction and deep sequencing for genomic studies. We also thank our Advanced Light Microscopy core (RRID: SCR_022526), Flow Cytometry core (RRID: SCR_021134), and Custom Antibody core (RRID: SCR_022799) for their service. We thank Dr. Elphege Pierre-Nora at UCSF for the generous gift of the CTCF-AID-GFP ESC line; and Dr. Diego Pasini (European Institute of Oncology IRCCS) for the generous gift of Pcgf2 antibody. This work was supported by the following grant to Z. Gao: R35GM133496 from NIGMS.

## Materials and Methods

### ESC culture, NPC, and neuronal differentiation

The ESC, NPC, and neuronal cell culture and differentiation methods were performed according to previously described protocols^38,39^. 10 cm tissue-culture-treated plates were coated with 0.1% gelatin and set for at least 15 minutes before aspiration. The plates were then seeded with γ-irradiated SNL feeder cells (ATCC), which are derived from mouse embryonic fibroblasts (MEFs) in the pre-warmed ESC medium, which consisted of DMEM (Corning, 10-017-CM) with 15% fetal bovine serum (FBS) (R&D Systems, S10250), 1 X non-essential amino acids (Gibco, 11140-050), 0.1mM β-mercaptoethanol (Fisher BioReagents, BP176-100), 0.5 X penicillin/streptomycin (Corning, 30-002-Cl), 1X sodium pyruvate (Corning, 25-000-Cl), 3.0 × 10^-3^ µg/mL LIF (Cayman Chemical, 32066), 1 μM PD0325901 (PD) (Cayman Chemical, 13034), and 3 μM Chir99021 (CH) (Cayman Chemical, 13122). The MEFs were allowed to settle and attach to the plates’ surface before plating ESCs on them. The co-cultures were incubated in an incubator set at 37°C with 5% CO_2_ content. The ESCs were differentiated into NPCs using the hanging drop method on 10 cm cell culture plates. 2,000 cells were suspended in 25 μL differentiation medium, which consisted of DMEM with 15% FBS, 1 X non-essential amino acids, 0.1 mM β-mercaptoethanol, 0.5 X penicillin/streptomycin, and 1 X sodium pyruvate. The droplet cultures were incubated in the incubator set at 37°C with 5% CO_2_ content. On day 2, the formed EBs were transferred from the droplets into suspension culture plates with 10 mL differentiation medium and left on an orbital shaker at low speed in the incubator. On day 3, some of the EBs were harvested for that timepoint. The rest of the EBs were cultured in differentiation medium either with 5 μM retinoic acid (RA) (Sigma, R2625) to induce the EBs’ differentiation into NPCs or without RA to maintain their EB identity. The old medium was replaced with a new differentiation medium on the subsequent days until day 8. Some of the NPCs and EBs were harvested on day 8 for that timepoint. To further differentiate the NPCs into neurons, the NPCs were dissociated and plated at a density of 1.5 × 10^5^/cm^2^ in N2 medium (DMEM/F12 medium with 3 mg/mL glucose, 3 mg/mL Albumax, 1/100 N2 supplement, 10 ng/mL bFGF, 50 U/mL pen/strep, and 1 mM L-glutamine). On day 9, the medium was changed. On day 10, the N2 medium was switched with N2/B27 medium (50% DMEM/F12 and 50% Neural Basal, 3 mg/mL Albumax, 1/200 N2 supplement, 1/100 B27 supplement, 50 U/mL penicillin/streptomycin, and 1 mM L-glutamine). The medium was refreshed for the next consecutive 2 days before being processed for downstream assays. Cells are passaged or harvested by covering the culture plates with Trypsin (Corning, 25-052-Cl).

### U2OS culture

U2OS cells (ATCC) were cultured in growth media consisting of DMEM (Corning, 10-017-CM), 10% FBS (R&D Systems, S11550), 10% newborn calf serum (NCS) (R&D Systems, S11250), and 0.5X penicillin/streptomycin (Corning, 30-002-Cl). Cells are plated on tissue culture plates and incubated in an incubator set at 37°C with 5% CO_2_ content. Cells are passaged or harvested by covering the culture plates with Trypsin.

### Knock-down using small interfering RNAs (siRNA)

U2OS cells were subjected to siRNA-mediated knock-down of either CTCF (siCTCF, Dharmacon, L-020165-00-0005) or RING1B (siRNF2, QIAGEN, S100095543). Each siRNA treatment was accompanied by a control (siControl, Dharmacon, D-001810-10-05). 20 µM of each siRNA was transfected into the cells using the Lipofectamine 2000 (Invitrogen, 11668019) reagent according to the manufacturer’s recommended protocol supplemented with OPTI-MEM medium (Gibco, 31985-070). The transfected cells were incubated at 37°C with 5% CO_2_ for 2 days before being processed for immunoprecipitation, immunoblotting, and immunofluorescence assay.

### CRISPR/Cas9-mediated gene editing

The CRISPR-Cas9-mediated gene editing in ESCs performed according to previously described method^80^. All genetic knock-out lines were done in E14 ESCs (ATCC). Cells were cultured as described above. The guide RNAs used to knock out Pcgf2, Pcgf4, Phc1, and truncation of the Phc1 SAM domain are listed in **Table S4**. The guide RNAs were cloned into the CRISPR/Cas9 plasmid system, PX458 (Addgene, Plasmid #48138). The plasmids containing the guide RNAs were transfected into E14 ESCs using the Lipofectamine 2000 reagent (Invitrogen, 11668019) as described by the manufacturer supplemented with OPTI-MEM medium (Gibco, 31985-070). The transfected cells were then sorted into single cells via flow cytometry for GFP-positive cells into 96-well plates. Live cells were sorted by flow cytometry under BSL-2 conditions using a FACSAria SORP (Becton Dickinson) instrument in Penn State College of Medicine’s Flow Cytometry core. The Pcgf2, Pcgf4, Phc1 knock-out, and Phc1 SAM domain truncation were confirmed by PCR using primers listed in **Table S4**. Further confirmation of the Pcgf2 and Pcgf4 deletions and Phc1 SAM domain truncation was done by Sanger sequencing (via GeneWiz) using the primers listed in **Table S4**. Prospective clones were also subjected to immunoblotting to confirm protein deletions and truncation.

### Immunoblotting

Whole-cell lysates were obtained from samples by using lysis buffer containing 1.5 mM MgCl_2_, 280 mM NaCl, 3 mM KCl, 0.15 mM EDTA, 15% EDTA,15 mM Tris-HCl pH 8.0, 0.02% IGEPAL, and protease inhibitors added fresh during the experiment (0.5 mM PMSF, 1 μg/mL Pepstatin A (Sigma-Aldrich, P5318), 1 μg/mL Leupeptin (Sigma-Aldrich, EI8), 1 μg/mL Aprotinin(Sigma-Aldrich, 616370). Protein concentration of samples was measured using the Bradford assay (Thermo Scientific, 1856209). The samples were loaded into SDS-PAGE gel to analyze the proteins extracted. The protein samples were transferred to nitrocellulose blot using the Trans-Blot Turbo system (BioRad). The blots were blocked in 5% milk and washed with 1X TBST. Refer to **Table S3** for the list of antibodies used for immunoblotting, immunoprecipitation, and ChIP-seq.

### Immunoprecipitation

Immunoprecipitation methods were adapted from the previously described protocol^18^. CTCF-AID-EGFP ESCs were first cultured as described in the ESC culture section above. When cells are at roughly 1.0 × 10^6^ density per sample, they were treated with 500 µM auxin in the form of indole-3-acetic acid (IAA) or DMSO at equal volume for 24 hours before being collected and processed for whole-cell lysate as described in the immunoblot section above. Protein lysates were incubated for 2 hours at 4°C with 10 µg of CTCF antibody in a volume of 400 μL Buffer A (10 mM Tris-HCl pH 8.0, 1.5 mM MgCl_2_, and 10 mM KCl) and 400 μL Buffer BN (20 mM Tris-HCl pH 8.0, 100 mM KCl, 0.2 mM EDTA, 20% glycerol, 0.5 mg/mL BSA, and 0.1% IGEPAL) supplemented with 0.5 mM PMSF, 1 μg/mL Pepstatin A, 1 μg/mL Leupeptin, and 1 μg/mL Aprotinin. 30 μL of protein G beads were then added and incubated at 4 °C overnight. Beads were then washed with Buffer BN 5 times, eluted with 100 μL glycine (0.1 M, pH 2.0), and neutralized by adding 6.5 μL Tris solution (1.5 M, pH 8.8). The eluates were mixed with 1X SDS sample buffer and analyzed by SDS-PAGE, followed by immunoblotting.

### Alkaline phosphatase (AP) assay

The assay was conducted according to the manufacturer’s instructions (Stemgent, 00-0055). The culture medium was aspirated, and the cells were washed with 2 mL of 1X PBST. 1 mL of Fix Solution was added, and the cells were incubated at room temperature (RT) for 5 minutes. The Fix Solution was then aspirated, and the fixed cells were washed with 2 mL of 1X PBST. The 1X PBST was aspirated, and 1.5 mL of freshly prepared AP Substrate Solution was added. The cells were incubated in the dark and wrapped with foil at room temperature for up to 15 minutes until the color changed. The reaction was stopped when the color turned bright to avoid non-specific staining by aspirating the AP Substrate Solution and washing the wells twice with 2 mL of 1X PBS. The cells were covered with 1X PBS or mounting medium to prevent drying, with AP expression resulting in a red or purple stain and the absence of AP expression resulting in no stain.

### MTT cell proliferation assay

The assay was conducted according to the manufacturer’s instructions (Invitrogen, 13154). The ESCs were cultured in 96-well plates. 10 μL of the 12-mM MTT stock solution was added to each well, and negative control was included by adding 10 μL of the MTT stock solution to 100 μL of medium alone. The wells were incubated at 37°C for 2 hours. All but 25 μL of the medium was aspirated from the wells. 50 μL of DMSO was added to each well, then pipetted up and down thoroughly to mix. The plates were incubated at 37°C for 10 minutes. Each sample was re-suspended by pipetting before the absorbance was read at 540 nm. This assay was done where measurements were taken every day for 4 days.

### Immunofluorescence assay

U2OS cells were cultured on tissue-culture-grade chamber slides (Thermo Fisher Scientific, 154526PK) for immunofluorescent assay. To prepare the chamber slides, they were coated with 0.1 mg/mL poly-L-ornithine and 0.1 mg/mL laminin. The chamber slides were washed with sterile 1X PBS before U2OS cells were plated and incubated at 37°C and 5% CO_2_. Once cells were ready to be processed, they were fixed with 4% formaldehyde for 15 minutes at RT. The chambers were then washed with 1X PBS. The cells were blocked and permeabilized with 5% BSA diluted in 1X PBS and 0.5% Triton-X100 solution for 1 hour. The chamber slides were washed with 1X PBS before incubating the samples in a primary antibody mix (1% BSA made in 1X PBS and 0.5% Triton X-100) for 1 hour at RT at dilution recommended by their respective manufacturers. After the chamber slides were washed with 1X PBS, the samples were then incubated in secondary fluorophore-conjugated antibodies mix for 1 hour at RT in the dark. The chamber slides were then washed with 1X PBS. Without leaving the samples completely dry, the chambers were removed from the slide before being stained with DAPI and mounted with a mounting solution. The samples were imaged with a fluorescence microscope. All quantifications calculated from images captured from immunofluorescent-stained samples were performed in ImageJ^81^.

### RNA-seq sample preparation

RNA-seq experiments were performed as previously described^63^. RNA was extracted using TRIzol following the manufacturer’s instructions (Invitrogen, 15596026). The RNA-seq mRNA library construction and sequencing procedures were performed by the Penn State College of Medicine Genomics Core using the TruSeq Stranded mRNA and Total RNA kit. Briefly, polyA RNA was purified from the total RNA using oligo (dT) beads. The extracted mRNA fraction was initially subjected to fragmentation, reverse transcription, end repair, 3′– end adenylation, and adaptor ligation. Then, the adaptor-ligated strands were subjected to PCR amplification and SPRISelect (Beckman Coulter) bead purification. The unique barcode sequences were incorporated into the adaptors for multiplexed high-throughput sequencing. The final product was assessed for size distribution and concentration using the BioAnalyzer High Sensitivity DNA Kit (Agilent, 5067-4626). Pooled libraries were diluted to 2 nM in EB buffer (Qiagen) and then denatured using the Illumina protocol. The denatured libraries were diluted to 10 pM by pre-chilled hybridization buffer and loaded onto a TruSeq Rapid flow cell on an Illumina Novaseq 6000 platform and run for 50 cycles using a single-read recipe according to the manufacturer’s instructions.

### RNA-seq analysis

Transcript abundances were estimated using the Kallisto program, with mm10 as the reference genome. The expected counts for each transcript were imported into R using the tximport package. Differentially expressed genes (DEGs) were identified using the Deseq2 package, with a fold-change larger than 2 and a p-adjusted value smaller than 0.01^82,83^. Upregulated and downregulated DEGs were used to perform Gene ontology (GO) analysis using the clusterprofiler package^84,85^. RNA-seq related plots were generated using ggplot2 and complexheatmap packages^86^. Neuro-ectoderm/meso-endo PRC1 loop gene list was generated by crossing the PRC1 loop gene list with the gene list acquired from GO term neurogenesis (GO:022008) and mesoderm development (GO:0007498) and endoderm development (GO:0007492) with manual curation. For RNA-seq visualization on IGV, the data processing was similar to that of ChIP-seq data.

### RT-qPCR

For the cDNA synthesis, 0.5 g of total extracted RNA was used, and the procedure was done using SuperScript III RT according to the manufacturer’s instructions (Invitrogen, 18080044). The cDNA samples were diluted 5 X before being used for the qPCR step.

qPCR was done according to the manufacturer’s instructions (Azura Genomics, AZ-2120). The reactions were performed and measured using the Biorad CFX Connect real-time PCR detection system. Refer to **Table S4** for the complete list of RT-qPCR primers used in this study.

### ChIP-seq sample preparation and sequencing

ChIP-seq samples were prepared as described previously^18^. Harvested cells were cross-linked with the fix solution (1% formaldehyde, 9 mM NaCl, 0.09 mM EDTA, 0.045 mM EGTA, 9 mM HEPES buffer pH 7.6) for 8 minutes. After washing and nuclei extraction, the samples were then resuspended in sonication buffer (0.5% N-Lauroyl Sarcosine, 1 mM EDTA, 0.5 mM EGTA, 10 mM Tris buffer pH 8.0, 0.5 mM PMSF, 1 µg/ml Pepstatin A, 1 µg/ml Leupeptin, and 1 µg/ml Aprotinin). The cross-linked nuclei pellets were sonicated using the Diagenode Bioruptor to about 200 bp after reverse cross-linking. To start the immunoprecipitation, the protein A/G beads were first washed with TE buffer and blocked for 1 hour at 4°C with 1 mg/mL BSA before being used to pre-clear chromatin samples. Each sample was immunoprecipitated using H2AK119ub1, Ring1b, and IgG antibodies in 3 X ChIP buffer (3% Triton X-100, 0.3% sodium deoxycholate, 3 mM EDTA, 0.5 mM PMSF, 1 µg/ml Pepstatin A, 1 µg/ml Leupeptin, and 1 µg/ml Aprotinin). The ChIP samples were then washed with RIPA buffer (50 mM HEPES pH7.6, 0.5M LiCl, 1mM EDTA, 1% IGEPAL, 0.7% DOC, 0.5 mM PMSF, 1 µg/ml Pepstatin A, 1 µg/ml Leupeptin, and 1 µg/ml Aprotinin) and 10% of each sample was loaded on SDS-PAGE for enrichment verification by immunoblotting. The remaining 90% of the ChIP samples were subjected to DNA extraction using the ethanol precipitation method with the PCI reagent (phenol: chloroform: isoamylalcohol) (Sigma-Aldrich, P3803). The ChIP-seq library and sequencing procedures were performed by the Penn State College of Medicine Genomics Core. The ChIP-seq library was constructed using the sparQ DNA Library Prep kit. Final libraries were subsequently sequenced using the NovaSeq platform (Illumina) as described^87^.

### ChIP-seq analysis

Raw FASTQ data was mapped using bwa against the mm10 mouse reference genome. Sam files were converted to bam files, followed by sorting and indexing using samtools. Peak calling was performed using MACS2 using aligned bam files^88^. For data visualization and downstream analysis, aligned bam files were converted to bigwig files via deepTools using bins per million mapped reads (BPM) normalization option^89^. Then, ChIP-seq data were visualized in bigwig format using an integrated genome browser (IGV) or Coolbox. For the heatmap and intensity plot, the selected ChIP-seq data and correlated genomic coordinate files were processed using the computeMatrix function, and plots were generated using the plotHeatmap function in deepTools^89^. Refer to **Table S5** for the full list of publicly available ChIP-seq, Hi-C, and HiChIP datasets used in this study.

### Hi-C sample preparation and library construction

The samples were prepared using the Arima-HiC+ kit (Arima Genomics, A510008) based on the manufacturer’s protocol with a few modifications. Briefly, cells were harvested based on the appropriate procedure to prepare samples for Hi-C and counted using a hemocytometer. For each sample, 2 million cells were cross-linked. This was done by resuspending the cells in 1 mL of 1 X PBS and cross-linked in 2% formaldehyde (Thermo Fisher Scientific, 28908) by inverting the tube 10 times before incubating at RT for 10 minutes. Glycine was added to each sample at the 0.25 M final concentration, and the tubes were inverted 10 times. The samples were then incubated at room temperature for 5 minutes and subsequently for 15 minutes on ice. The cells were centrifuged. The cell pellets were then washed with 1 X PBS and centrifuged again to remove the supernatant. The cross-linked cells were stored in the −80°C freezer before continuing with the subsequent steps. The library preparation procedures were conducted using the Arima Library Prep Module (Arima Genomics, A303011) based on the manufacturer’s instructions.

### Hi-C analysis

Raw FASTQ files were mapped against the mm10 mouse reference genome using the runHiC pipeline (http://dx.doi.org/10.5281/zenodo.55324). Briefly, the bwa aligner was used for alignments, and aligned reads were filtered for quality control and PCR duplication. Fragments were assembled and filtered to retain the fragments with at least two different restriction fragments to filter out self-ligated fragments. Last, the reads were binned at multiple resolutions to generate a contact matrix using cooler tools^90^. ICE-corrected Hi-C matrices were visualized using coolbox^91^. Hi-C data at 10KB resolution were used for look calling for each sample, which was achieved using pyHICCUPS from HiCPeaks packages^92^. Bedtools was used to compare the loops among samples with 2-mismatches and identify the overlap between Hi-C loops and ChIP-seq peaks of multiple proteins^93^. WT ESC Hi-C loops were crossed with ChIP-seq of Pcgf2 and Ring1b, and the loops with both occupied anchors were considered PRC1 loops. Aggregated plot analysis (APA) plots were generated using the APA analysis function from HiCPeaks^92^.

### Materials availability

Cell lines, plasmid constructs, and antibodies generated for this study are available upon request made to the lead contact, Zhonghua Gao, and processed via Penn State University material transfer agreement.

### Data availability

Raw and processed next-generation sequencing data generated in this study are deposited at the Gene Expression Omnibus (GEO) under the accession number GSE278946. All other ChIP-seq, Hi-C, and HiChIP datasets used from other studies are listed in **Table S5** with their corresponding identification or accession numbers. The bioinformatic analyses performed in this study utilized publicly available packages as described in the Materials and Methods section. Any additional information required to reproduce data reported in this article is available from the lead contact upon request.

### Author contributions

A.H., Z.Geng, T.L., Y.T., W.C., Q.W., and N.C. conducted the experiments; Z.Gao, F.Y., and Y.L. conceived the projects and designed the experiments; A.H., Z.Geng, and Z.Gao wrote the paper; All authors contributed to the discussion of the manuscript.

