## Supplemental Info for "PRC1 and CTCF-Mediated Transition from Poised to Active Chromatin Loops Drives Bivalent Gene Activation"

Figure S1

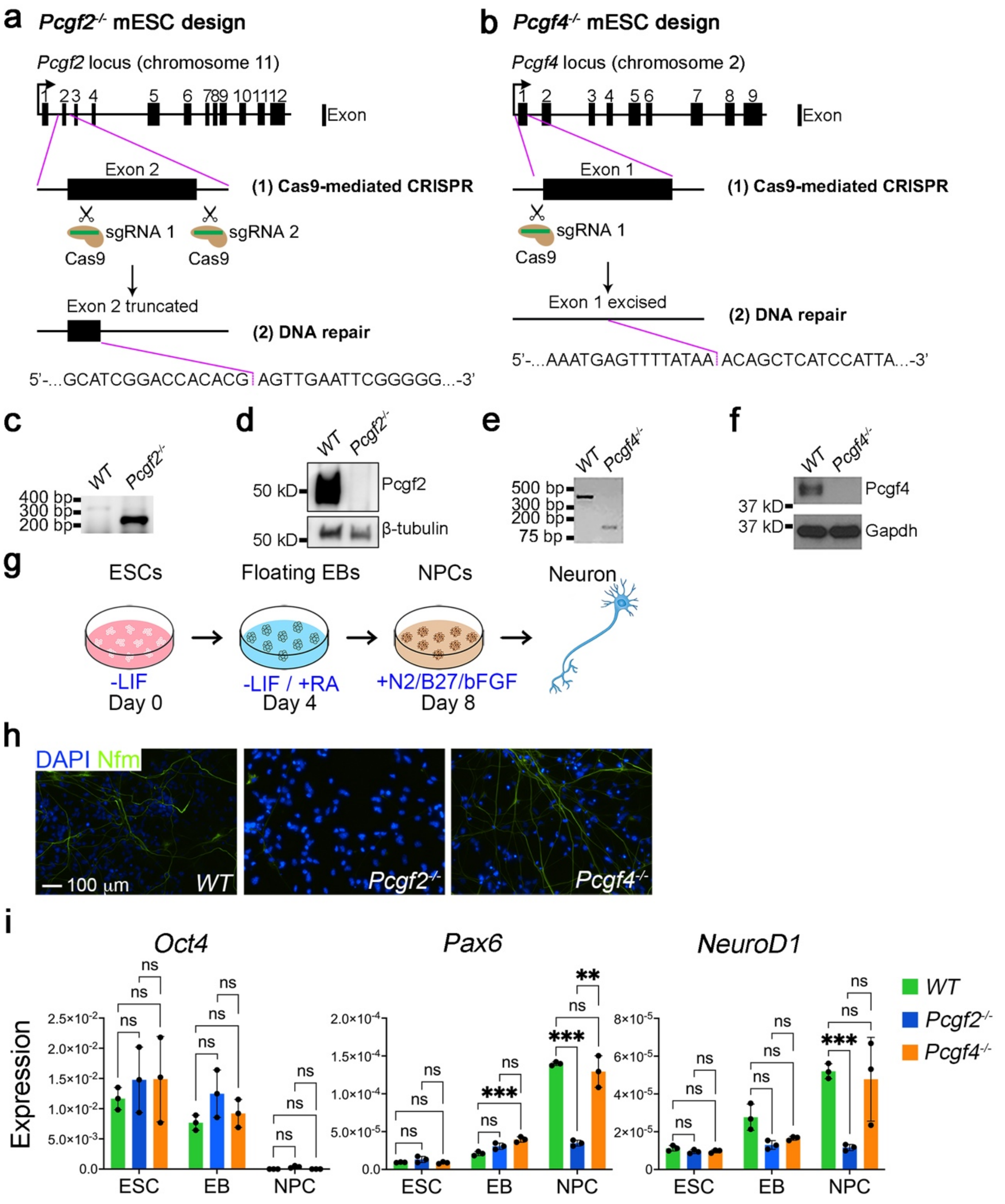


**Figure S1. Generation and phenotypic analysis of *Pcgf2*$^{-/-}$ and *Pcgf4*$^{-/-}$ ESC lines. a,** Schematic of CRISPR/Cas9-mediated deletion of Pcgf2 in ESCs. Bottom: Sanger sequencing validating the excision of Exon 2. **b,** Schematic of CRISPR/Cas9-mediated deletion of Pcgf4 in ESCs. Bottom: Sanger sequencing validating the excision of Exon 1. **c**, Gel electrophoresis of PCR products using primers targeting excision region in *WT* and *Pcgf2*$^{-/-}$ ESC clone. **d,** Immunoblotting of Pcgf2 in *WT* and *Pcgf2*$^{-/-}$ ESC clone. **e**, Gel electrophoresis of PCR products using primers targeting excision region in *WT* and *Pcgf4*$^{-}$

$^{/-}$ ESC clone. **f**, Immunoblotting of Pcgf4 in *WT* and *Pcgf4*$^{-/-}$ ESC clone. **g**, Schematic of NPC differentiation, as detailed in the Materials and Methods. **h**, Immunofluorescence staining of Neurofilament (Nfm), a neuronal marker, in neurons differentiated from *WT*, *Pcgf2*$^{-/-}$, *Pcgf4*$^{-/-}$ ESCs. **i**, RT-qPCR analysis of a pluripotency (*Oct4*) and NPC (*Pax6* and *NeuroD1*) markers in WT, *Pcgf2*$^{-/-}$, and *Pcgf4*$^{-/-}$ cells at the ESC, EB, and NPC stages, normalized to *18s rRNA*. Each data point represents three biological replicates. **$p<0.01$; ***$p<0.001$; n.s., not significant.

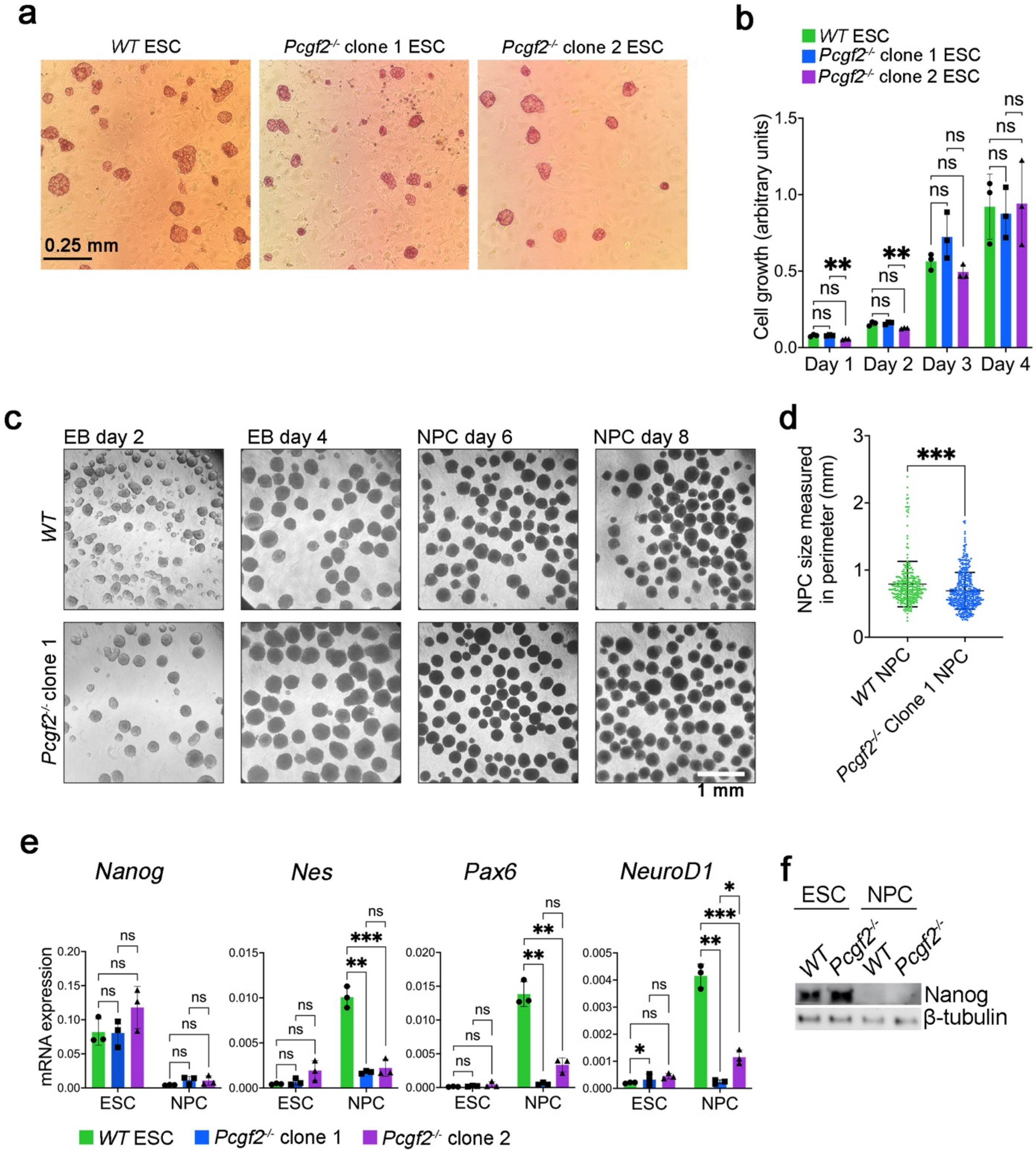


**Figure S2. Characterization of *Pcgf2*$^{-/-}$ ESC clones. a,** Alkaline phosphatase (AP) staining in *WT* and two *Pcgf2*$^{-/-}$ ESC clones co-cultured with mouse embryonic fibroblasts (MEF). Pluripotent cells stain purple. **b,** MTT cell proliferation assay in *WT* and two *Pcgf2*$^{-/-}$ ESC clones over four days. Each data point represents three biological replicates. **c**, Brightfield images of the *WT* and *Pcgf2*$^{-/-}$ clone 1 at EB and NPC stages during differentiation. **d**, Quantification of NPC size at day 8 in *WT* (n = 277) and *Pcgf2*$^{-/-}$ clone 1 (n = 416) based on cell perimeter. **e,** RT-qPCR of pluripotency (*Nanog*) and NPC (*Nes*, *Pax6*, and *NeuroD1*) markers in *WT* and two *Pcgf2*$^{-/-}$ clones at the ESC, EB, and NPC stages, normalized to *18s rRNA*. Each data point represents three biological replicates. **f**, Immunoblotting showing the Nanog protein level in *WT* and *Pcgf2*$^{-/-}$ clone 1 ESCs and NPCs. **p*<0.05; ***p*<0.01; ****p*<0.001; n.s., not significant.

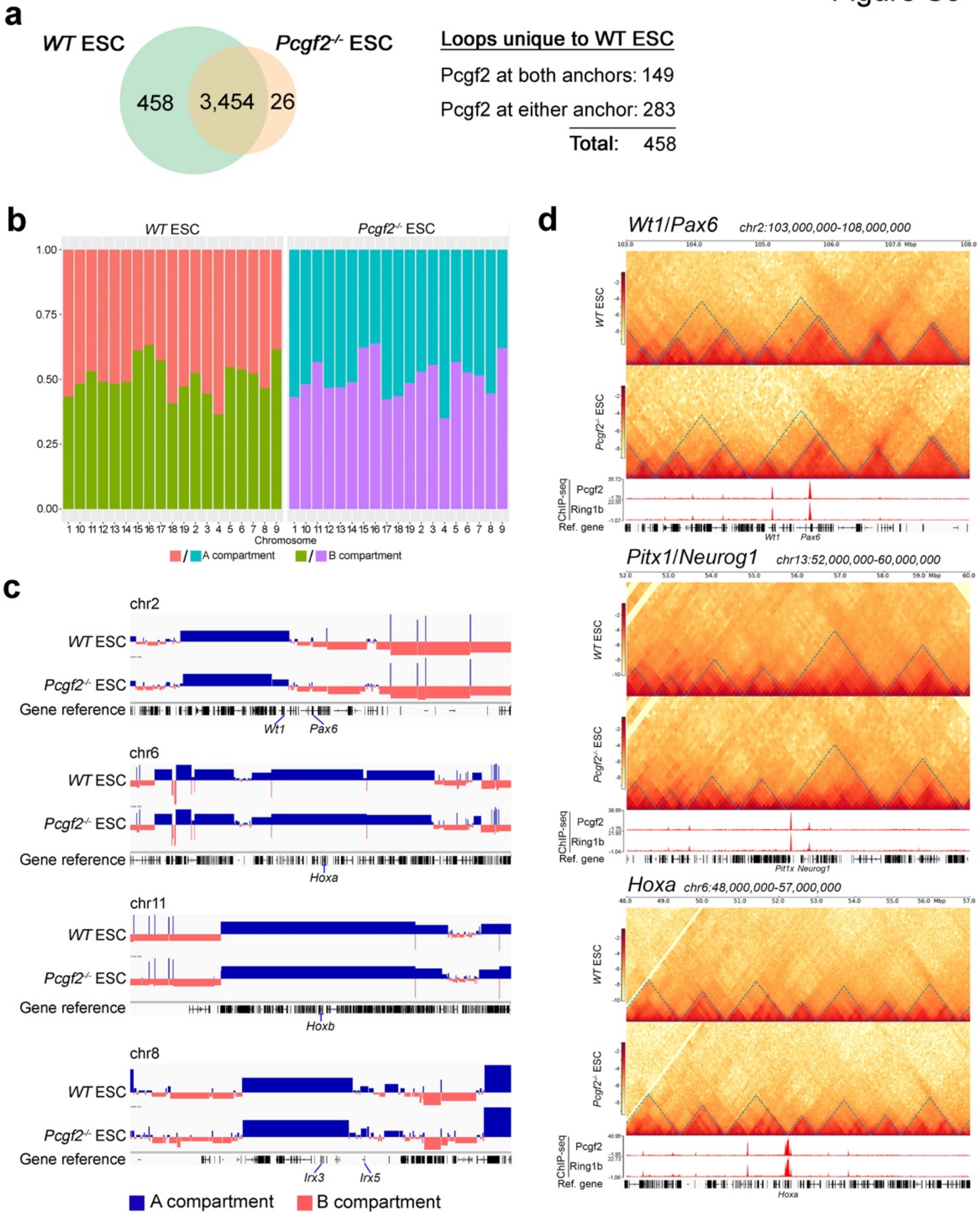


**Figure S3. Hi-C analysis in *WT* and *Pcgf2$^{-/-}$* ESCs. a**, Venn diagram showing the number of unique and shared chromatin loops in *WT* and *Pcgf2$^{-/-}$* ESCs. **b**, A/B compartment analysis for each autosomal chromosome in *WT* and *Pcgf2$^{-/-}$* ESCs derived from Hi-C data. **c**, A/B compartment plots at the *Wt1*/*Pax6*, *Hoxa*, *Hoxb*, and *Irx3*/*Irx5* loci. **d**, Contact frequency maps comparing *WT* and *Pcgf2$^{-/-}$* ESCs at *Wt1*/*Pax6*, *Pitx1*/*Neurog1*, and *Hoxa* loci. Topologically associated domains (TADs) are highlighted in dashed blue lines. The bottom panel shows ChIP-seq tracks for Ring1b and Pcgf2 in *WT* ESC.

Figure S4

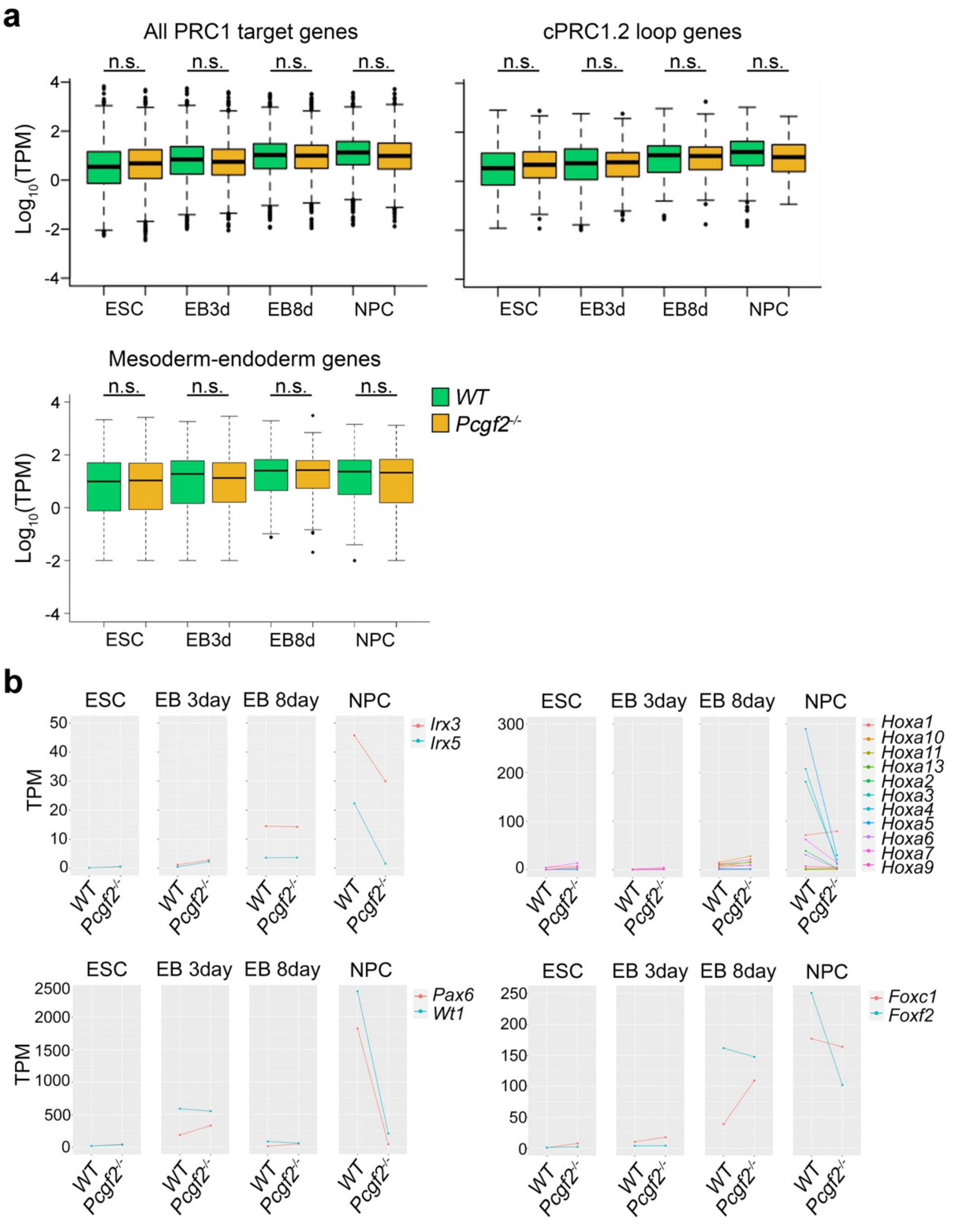


**Figure S4. Altered expression of cPRC1.2 loop target genes in *Pcgf2$^{-/-}$* cells. a**, Box plots show expression for all PRC1 target genes (5,334), total cPRC1.2 loop genes (454), and cPRC1.2 loop targeted meso-endodermal genes (155) in *WT* and *Pcgf2$^{-/-}$* cells at ESC, EB, and NPC stages. **b**, Line plots showing the transcript per million (TPM) values for genes within *Irx3*/*Irx5*, *Hoxa*, *Wt1*/*Pax6*, and *Foxf2*/*Foxc1* loci from RNA-seq data in *WT* and *Pcgf2$^{-/-}$* cells at ESC, EB, and NPC stages. n.s., not significant.

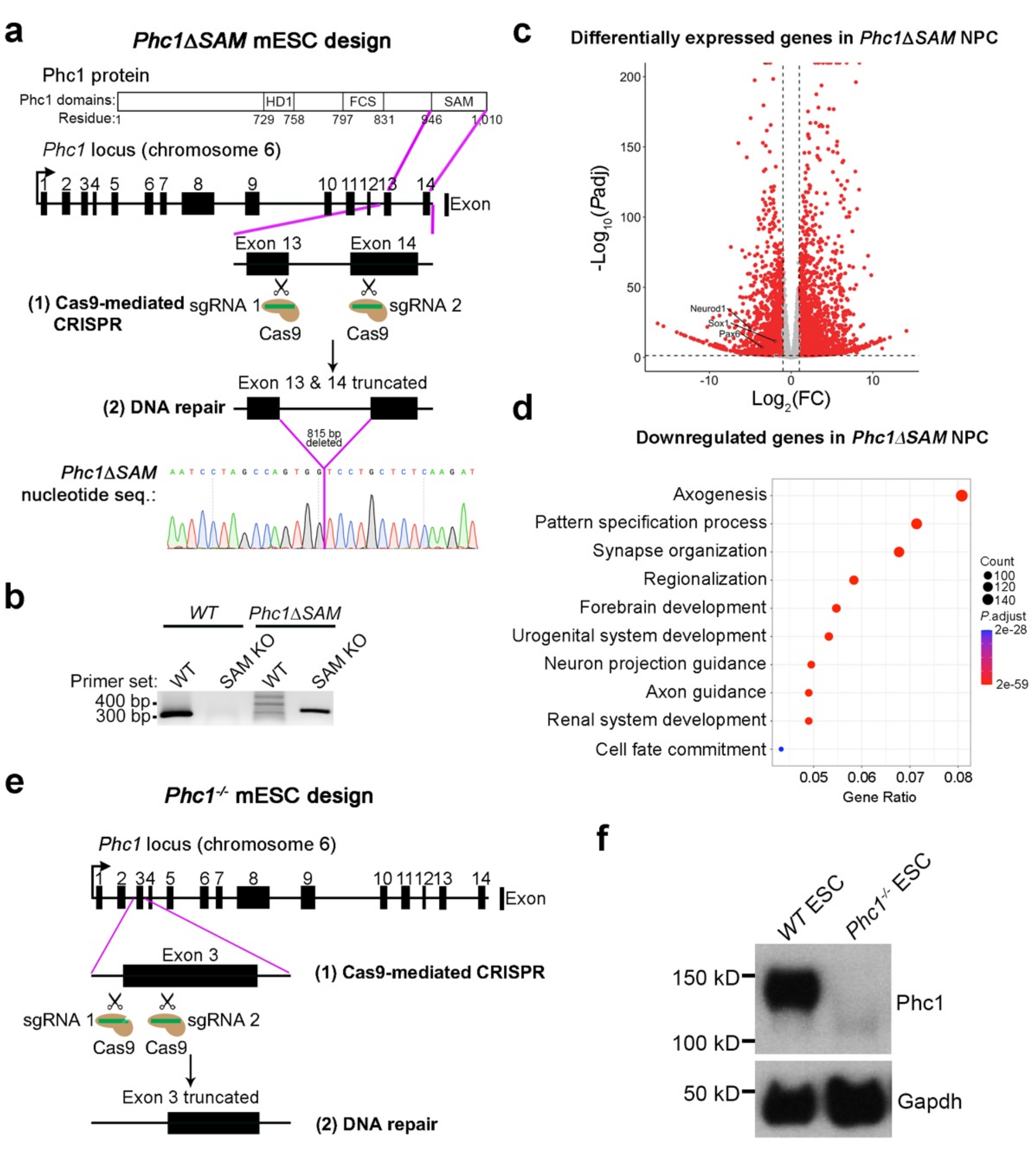


**Figure S5. SAM domain truncation and Phc1 deletion in ESCs. a,** Schematic of Phc1 SAM domain deletion (*Phc1ΔSAM*) in ESCs by CRISPR/Cas9-mediated gene editing. The SAM domain spans parts of exons 13 and 14. The bottom panel shows the Sanger sequencing result of a *Phc1ΔSAM* ESC clone with SAM domain excision. HD1: homology domain 1; FCS: FCS zinc-finger domain. **b**, PCR of a *WT* and *Phc1ΔSAM* ESC clone using WT and Δ*SAM*-specific primer set showing the successful deletion. **c**, Volcano plot of differentially expressed genes (DEGs) in *Phc1ΔSAM* NPCs compared to *WT* NPCs. Red dots denote significant DEGs, and gray dots denote insignificant DEGs. NPC markers such as *NeuroD1*, *Sox1*, and *Pax6* are indicated among the downregulated genes. **d**, GO analysis of downregulated genes in *Phc1ΔSAM* NPCs. Gene ratio (x-axis) indicates the proportion of genes in each GO term. **e**, Schematic showing the complete deletion of Phc1 (Exon 3) in ESCs. **f**, Immunoblotting showing the loss of Phc1 protein in *Phc1*$^{-/-}$ ESCs compared to *WT*.

Figure S6

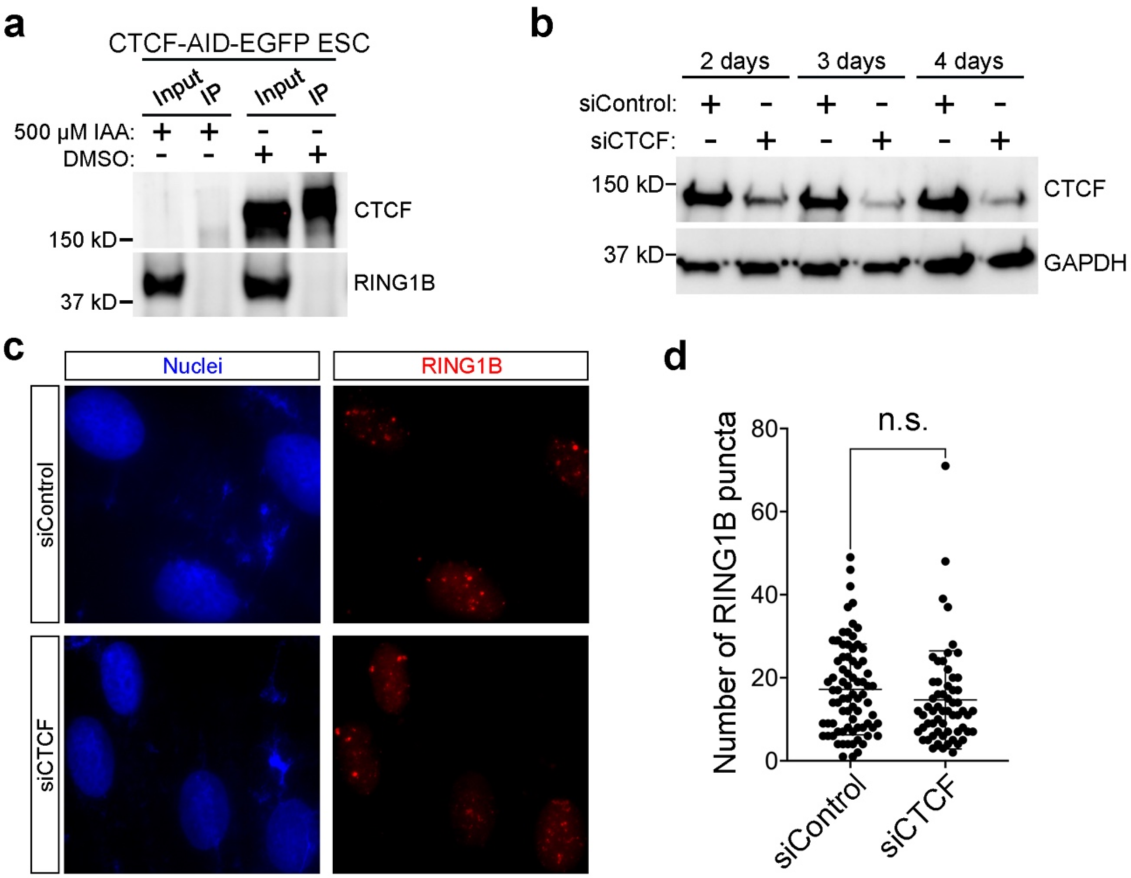


**Figure S6. CTCF is not required for PcG body formation. a**, Immunoprecipitation using CTCF antibody in CTCF-AID-EGFP ESCs with 500 μM indole-3-acetic acid (IAA) or DMSO treatment for 24 hours, followed by immunoblotting for CTCF and RING1B. **b**, Immunoblotting of CTCF in U2OS cells with CTCF knock-down (siCTCF) or control (siControl) over 2, 3, and 4 days. **c**, Immunofluorescent images of siControl and siCTCF U2OS cells stained for RING1B. **d**, Dot plots of PcG body count in siControl (n = 77) and siCTCF (n = 60) U2OS cells. n.s., not significant.

**Table S1.** Hi-C sequencing quality control parameters.

| **Parameter** | ***WT* ESC** | ***Pcgf2*$^{-/-}$ ESC** | ***Pcgf2*$^{-/-}$ NPC** |
|---|---|---|---|
| Total reads | 947053322 | 773871511 | 698183783 |
| Mapped reads | 661722797 | 539376595 | 499839643 |
| Mapping ratio | 661722797 / 947053322 = 0.6987 | 539376595 / 773871511 = 0.6970 | 499839643 / 698183783 = 0.7159 |
| Total reads after filtering | 397060244 | 192827488 | 406749595 |
| Inter-/trans-chromosomal reads | 39437227 | 19948992 | 80593419 |
| Intra-/cis-chromosomal reads | 357623017 | 172878496 | 326156176 |
| Intra-/cis-long range reads | 271779607 | 131376247 | 211555852 |

**Table S2.** List of cPRC1.2 loop target genes.

| Total cPRC1.2 loop target genes | | | | |
|---|---|---|---|---|
| *1700006H21Rik* | *Cngb1* | *Hes2* | *Mnat1* | *Six3os1* |
| *1700012A03Rik* | *Cntfr* | *Hes3* | *Mras* | *Slc22a18* |
| *1700018L02Rik* | *Col27a1* | *Hexa* | *Mrpl19* | *Slc28a1* |
| *1700022H16Rik* | *Col5a1* | *Hlx* | *Mrpl24* | *Slc38a3* |
| *1700039E22Rik* | *Col8a2* | *Hmx1* | *Mrpl33* | *Slc43a1* |
| *1700054K19Rik* | *Colgalt1* | *Hnf4g* | *Msantd2* | *Slc47a1* |
| *1700112H15Rik* | *Cpa2* | *Hotairm2* | *Msx1os* | *Slc47a2* |
| *2410002F23Rik* | *Crnde* | *Hottip* | *Mtor* | *Slc6a1* |
| *2610016A17Rik* | *Crtc3* | *Hoxa10* | *Mynn* | *Slc6a20a* |
| *2610027K06Rik* | *Ctsh* | *Hoxa11* | *Myo3b* | *Slc6a3* |
| *4921534H16Rik* | *Cyp26b1* | *Hoxa13* | *Myorg* | *Slc6a5* |
| *4931419H13Rik* | *D230030E09Rik* | *Hoxa2* | *Neurog1* | *Smarca5-ps* |
| *4933406D12Rik* | *D8Ertd738e* | *Hoxa5* | *Nkd1* | *Snai1* |
| *5730409E04Rik* | *Dbx1* | *Hoxaas2* | *Nkx2-1* | *Snx20* |
| *6430503K07Rik* | *Ddx4* | *Hoxb1* | *Nkx2-2os* | *Snx24* |
| *6430571L13Rik* | *Dgcr6* | *Hoxb13* | *Nkx2-9* | *Sorl1* |
| *9330179D12Rik* | *Dmrt2* | *Hoxb3* | *Nol4l* | *Sort1* |
| *9530036O11Rik* | *Dmrt3* | *Hoxb3os* | *Nop53* | *Sp3* |
| *A630075F10Rik* | *Dmrta2* | *Hoxb4* | *Noto* | *Sp8* |
| *A730035I17Rik* | *Dnajb12* | *Hoxb5* | *Nphp4* | *Sp9* |
| *A930004D18Rik* | *Dok7* | *Hoxb5os* | *Nr2e1* | *Spr* |
| *Abhd2* | *Dppa3* | *Hoxb6* | *Nr2f2* | *Spry1* |
| *Abhd8* | *Dpysl4* | *Hoxb7* | *Nr5a1* | *Srprb* |
| *Acan* | *Dusp2* | *Hoxb8* | *Nsun7* | *Stat5b* |
| *Acot7* | *Dysf* | *Hoxb9* | *Ntrk1* | *Steap1* |
| *Adamtsl3* | *Ech1* | *Hoxd10* | *Nxn* | *Steap2* |
| *Adcy7* | *Egfr* | *Hoxd11* | *Osbpl5* | *Stk32c* |
| *Adgre5* | *Elfn1* | *Hoxd13* | *Osr1* | *Stox2* |
| *Adpgk* | *En1* | *Hoxd3* | *Ostc* | *Stx18* |
| *Adra2c* | *En2* | *Hoxd8* | *Oxtr* | *Syndig1* |
| *Ago2* | *Espn* | *Hoxd9* | *Pafah2* | *Sypl2* |
| *Ago4* | *Etfbkmt* | *Hrh1* | *Parp11* | *Tal1* |
| *AI646519* | *Etnk2* | *Hrk* | *Parp16* | *Tbx18* |
| *Akap8l* | *Ets1* | *Hsd17b1* | *Paupar* | *Tbx2* |
| *Aldh4a1* | *Eva1a* | *Iffo2* | *Pax1* | *Tbx4* |
| *Alx4* | *Evx1* | *Igdcc3* | *Pax2* | *Tbx5* |
| *Amigo1* | *Evx1os* | *Igfbp4* | *Pax3* | *Tcp11* |
| *Ankle1* | *Evx2* | *Il27ra* | *Pax6* | *Tesc* |
| *Ankrd26* | *Fa2h* | *Immp1l* | *Pax7* | *Tet3* |
| *Ankrd63* | *Faim* | *Insyn1* | *Pax9* | *Tex45* |
| *Anks1* | *Fam43b* | *Iqgap1* | *Pde4dip* | *Tfap2a* |
| *Ano1* | *Fam50b* | *Irf8* | *Pdgfra* | *Tfap2b* |
| *Antxr2* | *Fam57a* | *Irx3* | *Peg10* | *Tfap2c* |

| | | | | |
|---|---|---|---|---|
| *Arhgef16* | *Fbln2* | *Irx5* | *Pglyrp2* | *Tfap2d* |
| *Arid1a* | *Fbln7* | *Irx6* | *Phox2b* | *Tfap2e* |
| *Arsb* | *Fbxo2* | *Itpripl1* | *Pisd-ps1* | *Thap3* |
| *Aspdh* | *Fgf15* | *Jak2* | *Pitx1* | *Tlr2* |
| *Atp5g3* | *Fgf20* | *Josd2* | *Pkdcc* | *Tlx1* |
| *Atp6v0d1* | *Fgf5* | *Jph2* | *Platr9* | *Tm9sf4* |
| *B830017H08Rik* | *Fgf8* | *Kazald1* | *Pls1* | *Tmco5* |
| *Bahd1* | *Fli1* | *Kcng3* | *Plxna4* | *Tmem115* |
| *Barhl2* | *Fosl2* | *Kcnj1* | *Pnmal2* | *Tmem167b* |
| *Bcar3* | *Foxa1* | *Kcnq1* | *Ppp1r9a* | *Tmem33* |
| *Bcas1os2* | *Foxa2* | *Kiss1* | *Prdm6* | *Tmem50b* |
| *Bcat1* | *Foxc1* | *Kiz* | *Prdm8* | *Tox2* |
| *Bcl11b* | *Foxd2* | *Klf4* | *Prickle1* | *Trappc9* |
| *Bmi1* | *Foxe3* | *Lbx1* | *Prkaca* | *Trim2* |
| *Bmp7* | *Foxf1* | *Ldhd* | *Ptgs1* | *Trp73* |
| *Brd7* | *Foxf2* | *Lef1* | *Pxdc1* | *Trpc1* |
| *C1qtnf12* | *Foxi2* | *Lhx2* | *Rab3a* | *Ttc29* |
| *Cacna1c* | *Foxj2* | *Lhx5* | *Rab6b* | *Ttll11* |
| *Cacna2d2* | *Foxo3* | *Limd1* | *Rasgrf1* | *Uchl1os* |
| *Cacna2d3* | *Foxp1* | *Lingo1* | *Resf1* | *Unc5b* |
| *Cacng6* | *Foxq1* | *Loxl1* | *Rex2* | *Uncx* |
| *Cacng7* | *Furin* | *Lrrc25* | *Rfxap* | *Urad* |
| *Calm3* | *Gad1* | *Lrrc36* | *Rgs12* | *Vangl1* |
| *Camk2n1* | *Gjd3* | *Lrtm2* | *Rn45s* | *Vax2os* |
| *Car7* | *Gm10390* | *Lypd6b* | *Rnf166* | *Vinac1* |
| *Casd1* | *Gm12505* | *Mad2l2* | *Rnf223* | *Wdr73* |
| *Casq2* | *Gm12830* | *Man1c1* | *Robo3* | *Whrn* |
| *Cbx4* | *Gm13212* | *Mcemp1* | *Rtn4rl2* | *Wnt7a* |
| *Cbx8* | *Gm14424* | *Mecom* | *Rxra* | *Wt1os* |
| *Ccdc141* | *Gm15050* | *Meis2* | *Sall1* | *Xrn2* |
| *Ccdc8* | *Gm15706* | *Mesp1* | *Samd7* | *Zdhhc2* |
| *Ccdc9b* | *Gm15825* | *Mest* | *Scamp1* | *Zeb2* |
| *Cd276* | *Gm20755* | *Micu1* | *Scube3* | *Zfp1* |
| *Cd82* | *Gm26688* | *Mir10b* | *Sdc2* | *Zfp120* |
| *Cdc20b* | *Gm29683* | *Mir12205* | *Sec61g* | *Zfp319* |
| *Cdkn1c* | *Gm38436* | *Mir125b-1* | *Selenok* | *Zfp467* |
| *Cdkn2c* | *Gm41289* | *Mir148a* | *Selenon* | *Zfp541* |
| *Cdx2* | *Gm6525* | *Mir1963* | *Sestd1* | *Zfp593* |
| *Cebpb* | *Gnas* | *Mir6367* | *Sfi1* | *Zfp638* |
| *Celrr* | *Gpn2* | *Mir688* | *Sfn* | *Zfp644* |
| *Celsr2* | *Gramd1b* | *Mir7020* | *Sfrp2* | *Zfp775* |
| *Cfap206* | *Grm8* | *Mir7225* | *Sfta3-ps* | *Zfp800* |
| *Chst14* | *Gsc2* | *Mir760* | *Sgce* | *Zfp831* |
| *Clmp* | *Gsx1* | *Mir8111* | *Sgpl1* | *Zfp992* |
| *Clptm1l* | *Gsx2* | *Mir9-3hg* | *Siae* | *Zfpm1* |

| | | | | |
|---|---|---|---|---|
| *Clrn3* | *Gtdc1* | *Mira* | *Six1* | *Zscan2* |
| *Clstn2* | *Gtpbp10* | *Mlkl* | *Six2* | *Zswim6* |
| *Cmtm3* | *Hdgfl3* | *Mmadhc* | *Six3* | |
| **Neuro-ectoderm cPRC1.2 loop genes** | | | | |
| *Adra2c* | *Foxa2* | *Hoxd3* | *Noto* | *Sall1* |
| *Barhl2* | *Foxc1* | *Irx3* | *Nphp4* | *Six1* |
| *Bmp7* | *Foxo3* | *Irx5* | *Ntrk1* | *Six2* |
| *Cacng7* | *Foxp1* | *Irx6* | *Paupar* | *Six3* |
| *Ccdc141* | *Gsx1* | *Lbx1* | *Pax1* | *Six3os1* |
| *Celsr2* | *Gsx2* | *Lef1* | *Pax2* | *Spr* |
| *Dbx1* | *Hdgfl3* | *Lhx2* | *Pax3* | *Tal1* |
| *Dmrta2* | *Hes2* | *Lhx5* | *Pax6* | *Tfap2c* |
| *Dpysl4* | *Hes3* | *Lingo1* | *Pax7* | *Tlx1* |
| *En1* | *Hexa* | *Lrtm2* | *Phox2b* | *Trp73* |
| *En2* | *Hmx1* | *Meis2* | *Pitx1* | *Uncx* |
| *Evx1* | *Hoxa2* | *Mesp1* | *Plxna4* | *Whrn* |
| *Fa2h* | *Hoxaas2* | *Mtor* | *Ppp1r9a* | *Wnt7a* |
| *Faim* | *Hoxb1* | *Myo3b* | *Prdm8* | *Zeb2* |
| *Fbxo2* | *Hoxb3* | *Nkx2-1* | *Rab3a* | |
| *Fgf20* | *Hoxb8* | *Nkx2-2os* | *Rasgrf1* | |
| *Foxa1* | *Hoxd10* | *Nkx2-9* | *Robo3* | |
| **Meso-endoderm cPRC1.2 loop genes** | | | | |
| *Acvr1* | *Dkk1* | *Hnf1b* | *Onecut1* | *Sox2* |
| *Acvr1b* | *Dll3* | *Hsbp1* | *Osr1* | *Sox7* |
| *Acvr2a* | *Dusp1* | *Htt* | *Otx2* | *Srf* |
| *Acvr2b* | *Dusp2* | *Inhba* | *Ovol1* | *Ssbp3* |
| *Angpt1* | *Dusp4* | *Itga5* | *Paf1* | *T* |
| *Angpt2* | *Dusp5* | *Itgav* | *Palb2* | *Tal1* |
| *Angpt4* | *Ecsit* | *Kdm6a* | *Pelo* | *Tbx1* |
| *Ankrd17* | *Eomes* | *Kdm6b* | *Pofut2* | *Tbx19* |
| *Arc* | *Epb41l5* | *Kif16b* | *Poglut1* | *Tbx3* |
| *Armc5* | *Epha2* | *Ldb1* | *Pou4f1* | *Tbx6* |
| *Axin1* | *Ets2* | *Lef1* | *Pou5f1* | *Tcf15* |
| *Bmp4* | *Etv2* | *Leo1* | *Ppp2ca* | *Tcf7l1* |
| *Bmp7* | *Exoc4* | *Lhx1* | *Prkaca* | *Tead1* |
| *Bmpr1a* | *Ext1* | *Macf1* | *Prkar1a* | *Tead2* |
| *Bmpr2* | *Ext2* | *Macroh2a1* | *Pthlh* | *Tgfb1* |
| *Bptf* | *Eya1* | *Med12* | *Pus7* | *Tlx2* |
| *Cdc73* | *Fgf8* | *Megf8* | *Rpl38* | *Tnrc6c* |
| *Cer1* | *Fgfr1* | *Mesd* | *Rps6ka6* | *Trp63* |
| *Cfc1* | *Fn1* | *Mesp1* | *Rtf1* | *Twsg1* |
| *Chrd* | *Foxc2* | *Mesp2* | *Sall1* | *Txnrd1* |
| *Churc1* | *Foxh1* | *Mixl1* | *Scx* | *Vegfa* |
| *Cited2* | *Gata4* | *Msgn1* | *Setd2* | *Vtn* |
| *Col12a1* | *Gata6* | *Nanog* | *Sfrp2* | *Wls* |

| | | | | |
|---|---|---|---|---|
| *Col5a1* | *Gdf1* | *Nckap1* | *Shh* | *Wnt11* |
| *Col5a2* | *Gdf3* | *Nf2* | *Smad1* | *Wnt3* |
| *Crb2* | *Gja1* | *Nkx2-5* | *Smad2* | *Wnt3a* |
| *Ctdnep1* | *Gpi1* | *Nodal* | *Smad3* | *Wnt5a* |
| *Ctnnb1* | *Hand1* | *Nog* | *Smad4* | *Wnt8a* |
| *Ctr9* | *Hdac1* | *Notch1* | *Smo* | *Yap1* |
| *Dab2* | *Hhex* | *Nr4a3* | *Snai1* | *Zfp36l1* |
| *Dand5* | *Hnf1a* | *Nup133* | *Sox17* | *Zic2* |

**Table S3.** List of antibodies used in this study.

| Antibody | Manufacturer | Catalog # | Application |
|---|---|---|---|
| Ring1b | Bethyl Laboratories | A302-869A | Immunoblot, immunofluorescence, ChIP-seq |
| Pcgf2 (Mel18) | Gift from Diego Passini | N/A | Immunoblot, immunofluorescence |
| Pcgf2 (Mel18) | Generated by Pennsylvania State University College of Medicine Custom Antibody core | N/A | Immunoblot |
| Rybp | Santa Cruz Biotechnology | SC374235 | Immunoblot |
| Phc1 | Cell Signaling Technology | 13768 | Immunoblot |
| Pcgf4 (Bmi1) | Bethyl Laboratories | A301-694A | Immunoblot |
| Gapdh | Invitrogen | MA5-15738 | Immunoblot |
| β-tubulin | Abcam | ab6046 | Immunoblot |
| Oct4 | Santa Cruz Biotechnology | sc-365509 | Immunoblot |
| Nanog | Invitrogen | PA1-41577 | Immunoblot |
| H2AK119ub1 | Cell Signaling Technology | 8240 | Immunoblot, ChIP-seq |
| Histone H3 | Bethyl Laboratories | A300-823A | Immunoblot |
| CTCF | BD Biosciences | BD612148 | Immunoblot, immunofluorescence, immunoprecipitation |
| Alexa Fluor 488 | Thermo Fisher Scientific | A28175 | Immunofluorescence |
| Alexa Fluor 568 | Thermo Fisher Scientific | A-11011 | Immunofluorescence |
| IgG | Cell Signaling Technology | 2729S | Immunoprecipitation |

**Table S4.** List of gRNAs, PCR and RT-qPCR primers used in this study.

| **gRNAs** | |
|---|---|
| **Target knock-out** | **gRNA sequence (5' → 3')** |
| *Pcgf2* | Guide 1: CACGGATTAAAATCACGGAG |
| | Guide 2: GTGAGTTGAATTCGGGGGTG |
| *Pcgf4* | Guide 1: TTATAAACCGCTCAGCATTC |
| | Guide 2: TACACAGTGTTCGCCTTCT |
| *Phc1* | Guide 1: ACTGCAGCGGCAACCTAATG |
| | Guide 2: GGGGGAGTTACTGCAAGGTT |
| *Phc1* (SAM domain) | Guide 1: TCCTAGCCAATGGAGCGTCG |
| | Guide 2: AGTGCCATGAACATCAAATT |

| **PCR primers** | |
|---|---|
| **Target knock-out** | **Primer sequence (5' → 3')** |
| *Pcgf2* | Forward: GGTGACACTTCCCAAAT |
| | Reverse: GAGCCAGAAGCTCACTCCTGG |
| *Pcgf4* | Forward: ATTCTGATCTCATTAGTAAATTCCTTT |
| | Reverse: CATAACTGTGAGGTTTACTTTCTTTT |
| *Phc1* | Forward: GAGTAGACTACTGAATCCAACATTG |
| | Reverse: CTGCAACCTTAATCTAAGAACCAC |
| *Phc1* (SAM domain) | Forward (Common): GCCGGGGATCAGATAATTCCA |
| | Reverse (WT): AATGTGGACCTGGTTCTGCT |
| | Reverse (KO): GTTCTGCCTCCAAACCCCAC |

| **Sanger sequencing primers** | |
|---|---|
| **Target knock-out** | **Primer sequence (5' → 3')** |
| *Pcgf2* | CCAGGAGTGAGCTTCTGGCTC |
| *Pcgf4* | ATTCTGATCTCATTAGTAAATTCCTTT |
| *Phc1* (SAM domain) | CCATCCACGCCAGAGTTACA |

| **RT-qPCR primers** | | |
|---|---|---|
| **Species** | **Gene** | **Sequence (5' → 3')** |
| Mouse | *18srRNA* | Forward: GCAATTATTCCCCATGAACG |
| | | Reverse: GGCCTCACTAAACCATCCAA |
| Mouse | *Nanog* | Forward: AGGCTTTGGAGACAGTGAGGTG |
| | | Reverse: TGGGTAAGGGTGTTCAAGCACT |
| Mouse | *Oct4* | Forward: AGATCACTCACATCGCCAATCA |
| | | Reverse: CGCCGGTTACAGAACCATACTC |
| Mouse | *Nes* | Forward: AGTGCCCAGTTCTAGTGGTGTCC |
| | | Reverse: CCTCTAAAATAGAGTGGTGAGGGTTG |
| Mouse | *NeuroD1* | Forward: CGAGTCATGAGTGCCCAGCTTA |
| | | Reverse: CCGGGAATAGTGAAACTGACGTG |

| | | |
|---|---|---|
| Mouse | *Pax6* | Forward: CTTGGGAAATCCGAGACAGA |
| | | Reverse: CTAGCCAGGTTGCGAAGAAC |

**Table S5.** List of ChIP-seq, Hi-ChIP, and Hi-C datasets used in this study

| **Sample** | **GEO/Encode ID** | **DOI of study article** |
|---|---|---|
| Hi-C mouse *WT* ESC | GSE96107 | 10.1016/j.cell.2017.09.043 |
| Hi-C mouse *WT* NPC | GSE96107 | 10.1016/j.cell.2017.09.043 |
| ChIP-seq Pcgf2 *WT* ESC | GSE122715 | 10.1016/j.molcel.2019.04.002 |
| ChIP-seq Rybp *WT* ESC | GSE42466 | 10.1016/j.celrep.2012.11.026 |
| ChIP-seq Cbx7 *WT* ESC | GSE42466 | 10.1016/j.celrep.2012.11.026 |
| ChIP-seq Cbx2 *WT* ESC | GSE89949 | 10.1016/j.molcel.2017.01.009 |
| ChIP-seq Phc1 *WT* ESC | GSE89949 | 10.1016/j.molcel.2017.01.009 |
| ChIP-seq H3K4me3 *WT* ESC | ENCSR000CGO | N/A |
| ChIP-seq H3K4me3 *WT* NPC | GSE96107 | 10.1016/j.cell.2017.09.043 |
| ChIP-seq H3K27me3 *WT* ESC | ENCSR059MBO | N/A |
| ChIP-seq CTCF *WT* ESC | GSE96107 | 10.1016/j.cell.2017.09.043 |
| ChIP-seq CTCF *WT* NPC | GSE96107 | 10.1016/j.cell.2017.09.043 |
| Hi-ChIP H3K4me3 *WT* ESC | GSE94452 | 10.1038/s41594-020-00539-5 |
| Hi-ChIP H3K4me3 *WT* NPC | GSE94452 | 10.1038/s41594-020-00539-5 |
| Hi-ChIP H3K4me3 Ctcf knockdown ESC | GSE94452 | 10.1038/s41594-020-00539-5 |
| Hi-ChIP H3K4me3 Ctcf knockdown NPC | GSE94452 | 10.1038/s41594-020-00539-5 |
| ChIP-seq H3K27ac *WT* ESC | ENCSR000CGQ | N/A |
| ChIP-seq H3K27ac *WT* NPC | GSE96107 | 10.1016/j.cell.2017.09.043 |
